# Principles of mRNA targeting via the Arabidopsis m^6^A-binding protein ECT2

**DOI:** 10.1101/2021.04.18.440342

**Authors:** Laura Arribas-Hernández, Sarah Rennie, Tino Köster, Carlotta Porcelli, Martin Lewinski, Dorothee Staiger, Robin Andersson, Peter Brodersen

**Affiliations:** University of Copenhagen, Copenhagen Plant Science Center, Ole Maaløes Vej 5, DK-2200 Copenhagen N; University of Copenhagen, Department of Biology, Ole Maaløes Vej 5, DK-2200 Copenhagen N; University of Bielefeld, Faculty of Biology, RNA Biology and Molecular Physiology, D-33615 Bielefeld

**Keywords:** m^6^A, ECT2, YTHDF, iCLIP, HyperTRIBE, plant, Arabidopsis.

## Abstract

Specific recognition of *N6*-methyladenosine (m^6^A) in mRNA by RNA-binding proteins containing a YT521-B homology (YTH) domain is important in eukaryotic gene regulation. The Arabidopsis YTH-domain protein ECT2 is thought to bind to mRNA at URU(m^6^A)Y sites, yet RR(m^6^A)CH is the canonical m^6^A consensus site in all eukaryotes and ECT2 functions require m^6^A binding activity. Here, we apply iCLIP (individual-nucleotide resolution cross-linking and immunoprecipitation) and HyperTRIBE (targets of RNA-binding proteins identified by editing) to define high-quality target sets of ECT2, and analyze the patterns of enriched sequence motifs around ECT2 crosslink sites. Our analyses show that ECT2 does in fact bind to RR(m^6^A)CH. Pyrimidine-rich motifs are enriched around, but not at m^6^A-sites, reflecting a preference for *N6*-adenosine methylation of RRACH/GGAU islands in pyrimidine-rich regions. Such motifs, particularly oligo-U and UNUNU upstream of m^6^A sites, are also implicated in ECT2 binding via its intrinsically disordered region (IDR). Finally, URUAY-type motifs are enriched at ECT2 crosslink sites, but their distinct properties suggest function as sites of competition between binding of ECT2 and as yet unidentified RNA-binding proteins. Our study provides coherence between genetic and molecular studies of m^6^A-YTH function in plants, and reveals new insight into the mode of RNA recognition by YTH-domain-containing proteins.

## Introduction

*N6*-methyladenosine (m^6^A) is the most abundant modified nucleotide in eukaryotic mRNA bodies. It is required for embryonic development and stem cell differentiation in several animals and plants (Zhong et al. 2008; Batista et al. 2014; Ping et al. 2014; Geula et al. 2015; Zhang et al. 2017) and for the control of the meiotic program in yeast (Shah and Clancy 1992; Clancy et al. 2002; Agarwala et al. 2012). Most *N6*-adenosine methylation of mRNA is catalyzed in the nucleus (Salditt-Georgieff et al. 1976; Ke et al. 2017; Huang et al. 2019) by a highly conserved, multimeric methylase (the m^6^A “writer”) (Balacco and Soller 2019) whose catalytic core consists of the heterodimer METTL3/METTL14 (MTA/MTB in plants) (Bokar et al. 1997; Zhong et al. 2008; Liu et al. 2014). In addition, a number of highly conserved proteins is required for *N6*-methylation *in vivo* (Balacco and Soller 2019). The strong conservation of these core factors suggests that the biochemical basis of *N6*-adenosine methylation is common in eukaryotes and indeed, m^6^A occurs in the consensus site RR(m^6^A)CH (R=G/A, H=A/C/U), primarily in 3’-UTRs in animals (insects, mammals and fish), plants (maize and Arabidopsis) and fungi (yeast) that possess the canonical METTL3/METTL14 methyltransferase (Dominissini et al. 2012; Meyer et al. 2012; Schwartz et al. 2013; Luo et al. 2014; Lence et al. 2016; Zhao et al. 2017; Li et al. 2019; Miao et al. 2019; Parker et al. 2020). Conversely, the characteristic motif and gene-body location is not detected in organisms that lack METTL3/METTL14 homologs, such as the nematode *Caenorhabditis elegans* (Sendinc et al. 2020) and bacteria (Deng et al. 2015).

m^6^A may impact mRNA function by different mechanisms, including the creation of binding sites for reader proteins that specifically recognize m^6^A in mRNA (Dominissini et al. 2012; Fu et al. 2014; Meyer and Jaffrey 2014). The best understood class of readers contains a so-called YT521-B homology (YTH) domain (Stoilov et al. 2002) of which two phylogenetic groups, YTHDF and YTHDC, have been defined (Patil et al. 2018; Balacco and Soller 2019). The YTH domain harbors a hydrophobic methyl-binding pocket that increases the affinity of m^6^A-containing RNA by more than 10-fold compared to unmethylated RNA (Li et al. 2014b; Luo and Tong 2014; Theler et al. 2014; Xu et al. 2014; Zhu et al. 2014). Apart from interactions with the methylated adenosine and the purine at the -1 position, YTH-RNA domain interactions mostly involve the sugar-phosphate backbone of the RNA (Luo and Tong 2014; Theler et al. 2014; Xu et al. 2014). That is consistent with only mild reductions in the binding affinity of the YTH domain of human YTHDC1 upon substitution of nucleotides -2, +1 and +3 that abrogate the canonical RR(m^6^A)CH motif (Xu et al. 2014), and poor sequence specificity of RNA binding by isolated YTH domains of human YTHDF1, YTHDF2 and YTHDC1 (Arguello et al. 2019). Thus, the methyltransferase complex gives the sequence specificity, while YTH domain proteins may bind to m^6^A-containing RNA regardless of the identity of the immediately adjacent nucleotides.

YTHDF proteins are typically cytoplasmic and consist of a long N-terminal intrinsically disordered region (IDR) followed by the globular YTH domain (Patil et al. 2018). Because the affinity of isolated YTH domains for m^6^A-containing RNA is modest, typically with dissociation constants on the order of 0.1-1 μM (Li et al. 2014b; Luo and Tong 2014; Theler et al. 2014; Xu et al. 2014; Zhu et al. 2014), it has been suggested that the IDR may participate in RNA binding (Patil et al. 2018). Nonetheless, the clearest evidence for functions of the IDRs in YTHDF proteins reported thus far includes direct interactions with effectors such as the CCR4-NOT complex in mammalian cells (Du et al. 2016), and the ability to cause liquid-liquid phase transition when sufficiently high local concentrations are reached (Arribas-Hernández et al. 2018; Gao et al. 2019; Ries et al. 2019; Fu and Zhuang 2020; Wang et al. 2020).

The YTHDF family comprises 11 proteins in *Arabidopsis* that are referred to as <u>E</u>VOLUTIONARILY <u>C</u>ONSERVED C-<u>T</u>ERMINAL REGION1-11 (ECT1-11) (Li et al. 2014a; Scutenaire et al. 2018). ECT2, ECT3 and ECT4 are expressed in rapidly dividing cells of root, leaf and flower primordia and genetic analyses have revealed their general importance in development. Simultaneous inactivation of *ECT2* and *ECT3* causes slow organogenesis and abnormal morphology of leaves, roots, stems, flowers, and fruits, and these defects are generally enhanced by additional mutation of *ECT4* (Arribas-Hernández et al. 2018; Arribas-Hernández et al. 2020). Importantly, the biological functions of ECT2/3/4 described thus far are shared with those of m^6^A writer components and, where tested, have been shown to depend on intact m^6^A-binding pockets, strongly suggesting that the basis for the observed phenotypes in *ect2/3/4* mutants is defective regulation of m^6^A-modified mRNA targets (Bodi et al. 2012; Shen et al. 2016; Růžička et al. 2017; Arribas-Hernández et al. 2018; Scutenaire et al. 2018; Wei et al. 2018; Arribas-Hernández et al. 2020). Despite this progress in identifying biological functions of plant m^6^A-YTHDF axes, a number of fundamental questions regarding their molecular basis remains unanswered. For example, it is unclear whether sequence determinants in addition to m^6^A are important for mRNA target association of ECT proteins *in vivo*, the mRNA targets of ECT2/3/4 responsible for the developmental delay of *ect2*/*ect3/*(*ect4*) mutants have not been identified, and it is not clear what the effects of ECT2/ECT3/ECT4 binding to them may be (Arribas-Hernández and Brodersen 2020). Clearly, robust identification of the mRNA targets directly bound by ECT proteins is key to obtain satisfactory answers to all of these questions. Towards that goal, formaldehyde crosslinking and immunoprecipitation (FA-CLIP) was used to identify mRNA targets of ECT2 (Wei et al. 2018). Nonetheless, because formaldehyde, in contrast to UV illumination, generates both protein-protein and protein-RNA crosslinks, it is not an ideal choice for identification of mRNAs bound directly by a protein of interest (see Arribas-Hernández and Brodersen (2020) for a discussion). In particular, this problem concerns the unexpected conclusion that ECT2 binds to the ‘plant-specific consensus motif’ URU(m^6^A)Y (Y=U/C), not RR(m^6^A)CH (Wei et al. 2018). Thus, the field of gene regulation via m^6^A-YTHDF modules in plants is in a state of confusion: On the one hand, m^6^A mapping (Luo et al. 2014; Wan et al. 2015; Shen et al. 2016; Duan et al. 2017; Anderson et al. 2018; Miao et al. 2019; Parker et al. 2020) and phenotypes of mutants defective in m^6^A writing (Bodi et al. 2012; Shen et al. 2016; Růžička et al. 2017) or m^6^A-binding of ECT2/ECT3/ECT4 (Arribas-Hernández et al. 2018; Arribas-Hernández et al. 2020) suggest that these YTHDF proteins should act via recognition of m^6^A in the RR<u>A</u>CH context. On the other hand, the only attempt at a mechanistic understanding of ECT2 function via mRNA target identification concluded that ECT2 binds to a sequence element different from RRACH (Wei et al. 2018). To complicate matters further, a number of motifs including not only URUAY, but also UGUAMM (M=A/C), UGWAMH (W=A/U), UGUAWA and GGAU have been reported to be enriched around m^6^A sites in *Arabidopsis* and other plant species (Li et al. 2014c; Anderson et al. 2018; Miao et al. 2019; Zhang et al. 2019; Zhou et al. 2019), but it remains unclear whether the adenosines in such motifs are methylated *in vivo*. Alternatively, these sequence contexts may play a role in guiding m^6^A deposition or ECT recognition nearby, either directly by ECT interaction or indirectly via additional RNA binding proteins assisting or competing with ECT binding.

To clarify principles underlying mRNA recognition by ECT2, we undertook rigorous analysis of its mRNA binding sites using two orthogonal methods, the proximity-labeling method HyperTRIBE (targets of <u>R</u>NA binding proteins identified <u>b</u>y <u>e</u>diting) (McMahon et al. 2016; Xu et al. 2018), and iCLIP (individual nucleotide resolution <u>c</u>rosslinking and immuno<u>p</u>recipitation) (König et al. 2010). This resulted in identification of high-quality target sets as judged by mutual overlaps and by overlaps with previously reported m^6^A maps from plants at a similar developmental stage (Shen et al. 2016; Parker et al. 2020). Relying on this high-quality target set, we used the position information inherent to iCLIP and a single-nucleotide resolution m^6^A dataset (Parker et al. 2020) to establish six properties of m^6^A-containing mRNA and mRNA targeting by ECT2. (1) RRACH and its variant DR<u>A</u>CH (D=R/U) are unequivocally the most highly enriched motifs at m^6^A sites in Arabidopsis. (2) ECT2 binds to m^6^A sites in the canonical RR<u>A</u>CH context, as ECT2 crosslinking sites are preferentially found immediately 5’ to m^6^A sites, and RR<u>A</u>CH is enriched immediately 3’ to ECT2 crosslinking sites. (3) GG<u>A</u>U is a minor m^6^A consensus site in plants. (4) U- and U/C-rich motifs are enriched around, but not at, m^6^A sites, and, together with RR<u>A</u>CH and GG<u>A</u>U, constitute core elements that distinguish m^6^A-containing 3’-UTRs from non-m^6^A-containing 3’-UTRs in plants. (5) The IDR of ECT2 participates in RNA binding as it crosslinks to target mRNAs at U-rich elements highly abundant upstream of m^6^A-sites. (6) Although URUAY, URURU and similar motifs may crosslink to ECT2, their presence in m^6^A-containing mRNA disfavors ECT2 binding, consistent with those motifs acting predominantly as sites of interaction for RNA-binding proteins that may compete with ECT2.

## Results

### ADARcd fusions to ECT2 are functional in vivo

HyperTRIBE uses fusion of RNA binding proteins to the hyperactive E488Q mutant of the catalytic domain of the *Drosophila melanogaster* adenosine deaminase acting on RNA (*Dm*ADAR^E488Q^cd) (Kuttan and Bass 2012) to achieve proximity labeling *in vivo* (McMahon et al. 2016; Xu et al. 2018). Targets are identified as those mRNAs that contain adenosine-inosine sites significantly more highly edited than background controls, measured as A-G changes upon reverse transcription and sequencing. To develop material suitable for ECT2 HyperTRIBE, we expressed *AtECT2pro:AtECT2-FLAG-DmADAR^E488Q^cd-AtECT2ter* (henceforth “*ECT2-FLAG-ADAR*”) in the single *ect2-1* and triple *ect2-1/ect3-1/ect4-2* (*te234*) knockout backgrounds (Arribas-Hernández et al. 2018; Arribas-Hernández et al. 2020). We identified lines exhibiting nearly complete rescue of *te234* mutant seedling phenotypes, indicating that the fusion protein was functional (*Figure 1A*). We then used the expression level in complementing lines as a criterion to select lines in the *ect2-1* single mutant background, for which no easily scorable phenotype has been described (*Figure 1— figure supplement 1A*). Lines expressing free *Dm*ADAR^E488Q^cd under the control of the endogenous *ECT2* promoter (*AtECT2pro:FLAG-DmADAR^E488Q^cd-AtECT2ter;* henceforth *FLAG-ADAR*) at levels similar to or higher than those of the fusion lines (*Figure 1—figure supplement 1A,B*) were used to control for background editing after verification that *FLAG-ADAR* expression did not result in phenotypic abnormalities in Col-0 WT plants (*Figure 1A*).

**Figure 1.**
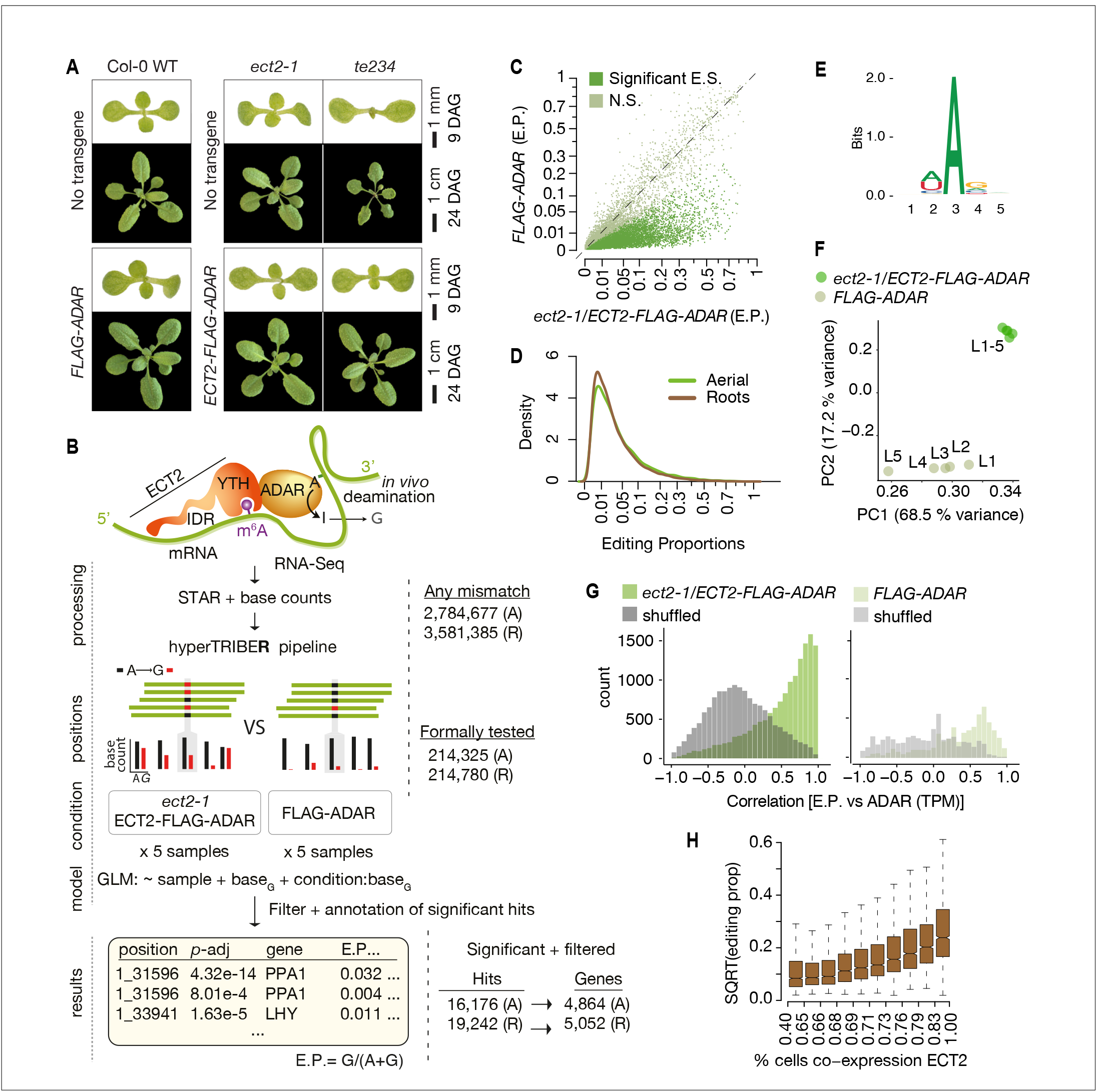
*Drosophila* ADARcd fused to ECT2 can edit target mRNAs *in vivo* in plants. **(A)** Phenotypes of wild type, *ect2-1* and *te234* mutants with (lower panels) or without (upper panels) *ECT2-FLAG-ADAR* or *FLAG-ADAR* transgenes, at 9 or 24 days after germination (DAG). **(B)** Experimental design for ECT2-HyperTRIBE (ECT2-HT) target identification and hyperTRIBE**R** pipeline. Nucleotide base counts quantified from mapped RNA-seq libraries were passed into the HyperTRIBE**R** pipeline to call significant editing sites, which were further filtered and annotated. The number of sites in either aerial (A, dissected apices) or root (R, root tips) tissues considered at each stage of the analysis is indicated. GLM, generalized linear model; E.P., editing proportion. **(C)** Scatterplot of the editing proportions of potential and significant editing sites (E.S.) in aerial tissues of *ect2-1/ECT2-FLAG-ADAR* lines compared to the *FLAG-ADAR* controls. Significant sites are highlighted in vivid green. N.S., not significant. **(D)** Consensus motif identified at significant editing sites in aerial tissues of *ect2-1/ECT2-FLAG-ADAR* lines. **(E)** Principal component analysis of editing proportions at significant editing sites in samples with aerial tissue. **(F)** Density of editing proportions for significant editing sites in aerial tissues and roots of *ect2-1/ECT2-FLAG-ADAR* lines. **(G)** Distribution of the correlations between editing proportions and ADAR expression (TPM) for significant editing sites in aerial tissues of either *ect2-1/ECT2-FLAG-ADAR* or *FLAG-ADAR* lines. Background correlations are based on randomly shuffling ADAR expression for each site. **(H)** Boxplots showing the mean editing proportions as a function of the proportion of cells co-expressing *ECT2*, calculated based on single cell RNA-seq in roots (Denyer et al., 2019). For panels C, E, and G, comparable analyses in both aerial and root tissues are shown in the figure supplement 1. **Figure supplement 1.** *Drosophila* ADARcd fused to ECT2 can edit target mRNAs *in vivo i*n plants (extended data, aerial and root tissues).

### The ECT2-ADARcd fusion imparts adenosine-to-inosine editing of target mRNAs in planta

To identify ECT2 HyperTRIBE targets (HT-targets), we sequenced mRNA from dissected root tips and shoot apices of 10-day-old seedlings of *ect2-1*/*ECT2-FLAG-ADAR* and *FLAG-ADAR* transgenic lines, using five independent lines of each type as biological replicates to prevent line-specific artifacts. Next, we generated nucleotide base counts for all positions with at least one mismatch across the full set of samples of mapped reads (*Figure 1B*), resulting in a raw list of potential editing positions. This revealed that the amount of editing was clearly higher in the lines expressing the ECT2-FLAG-ADAR fusion protein than in the negative control lines (*Figure 1C, Figure 1—figure supplement 1C*). To identify positions with significantly higher editing rates in ECT2-FLAG-ADAR lines compared to controls, we developed a new approach to detect differential editing (*Figure 1B*) that will be described in detail in a subsequent report. Briefly, the hyperTRIBE**R** (https://github.com/sarah-ku/hyperTRIBER) method of detecting differential editing exploits the powerful statistical capabilities of a method originally designed to detect differential exon usage (Anders et al. 2012). It efficiently takes replicates and possible differences in expression into account, resulting in high power to detect sites despite the generally low editing proportions that we found in our data (*Figure 1D*). As expected, the tendency towards higher editing proportions in fusion lines compared to controls was even more pronounced after filtering non-significantly edited sites (*Figure 1C, Figure 1—figure supplement 1C*). Three additional properties of the resulting editing sites indicate that they are the result of ADARcd activity guided by its fusion to ECT2. First, the vast majority of significant hits corresponded to A-to-G transitions (*Figure 1—figure supplement 1D*). Second, the consensus motif at the edited sites matched the sequence preference of *Dm*ADAR^E488Q^cd (5’ and 3’ nearest-neighbor preference of U>A>C>G and G>C>A∼U, respectively (Xu et al. 2018)) (*Figure 1E, Figure 1—figure supplement 1E*), with highly edited sites more closely matching the optimal sequence context than lowly edited ones (*Figure 1—figure supplement 1F*). Third, principal component analysis of editing proportions at significant sites over the different lines clearly separated the ECT2-FLAG-ADAR fusion lines from the control lines (*Figure 1F, Figure 1— figure supplement 1G*). Application of subsequent filtering steps, including removal of non-(A-to-G) mismatches and of potential line-specific single-nucleotide variants (see *Methods*), resulted in a final list of 16,176 edited sites for aerial tissues and 19,242 for roots, corresponding to 4,864 and 5,052 genes (ECT2 HT-targets), respectively (*Figure 1B*, *Supplementary file 1*). In both cases, this represents 27% of all expressed genes. We note that the editing proportions were generally low (*Figure 1D*) compared to previous work in Drosophila (Xu et al. 2018), perhaps in part due to the limited number of cells that express ECT2 (Arribas-Hernández et al. 2018; Arribas-Hernández et al. 2020). Indeed, the *ADAR* expression level (TPMs) correlated strongly with editing proportions among *ECT2-FLAG-ADAR* lines (*Figure 1G, Figure 1—figure supplement 1H*), and editing proportions were higher for target mRNAs that are co-expressed with *ECT2* in a large percentage of cells according to single-cell RNA-seq (Denyer et al. 2019) (*Figure 1H*), lending further support to the conclusion that the observed editing is ADAR-specific and driven to target mRNAs by ECT2. Hence, HyperTRIBE can be used to identify targets of RNA binding proteins *in planta*.

### HyperTRIBE is highly sensitive and identifies primarily m^6^A-containing transcripts as ECT2 targets

To evaluate the properties of ECT2 HT-targets, we first noted that most of them were common between root and aerial tissues (*Figure 2A*), as expected given the recurrent function of ECT2 in stimulating cell division in all organ primordia (Arribas-Hernández et al. 2020). In agreement with this result, most of the targets specific to root or aerial tissue were simply preferentially expressed in either tissue (*Figure 2B*). Moreover, the significant editing sites in roots and aerial tissues had a considerable overlap (*Figure 2A*), and their editing proportions were similar in the two tissues (*Figure 2C*). Of most importance, we observed a large overlap between the ECT2 HT-targets and m^6^A-containing transcripts mapped by different methods in seedlings (Shen et al. 2016; Parker et al. 2020), as more than 76% of ECT2 HT-targets had m^6^A support by either study (*Figure 2D*). These results validate our HyperTRIBE experimental setup and data analysis, and confirm that ECT2 binds predominantly to m^6^A-containing transcripts *in vivo*.

**Figure 2.**
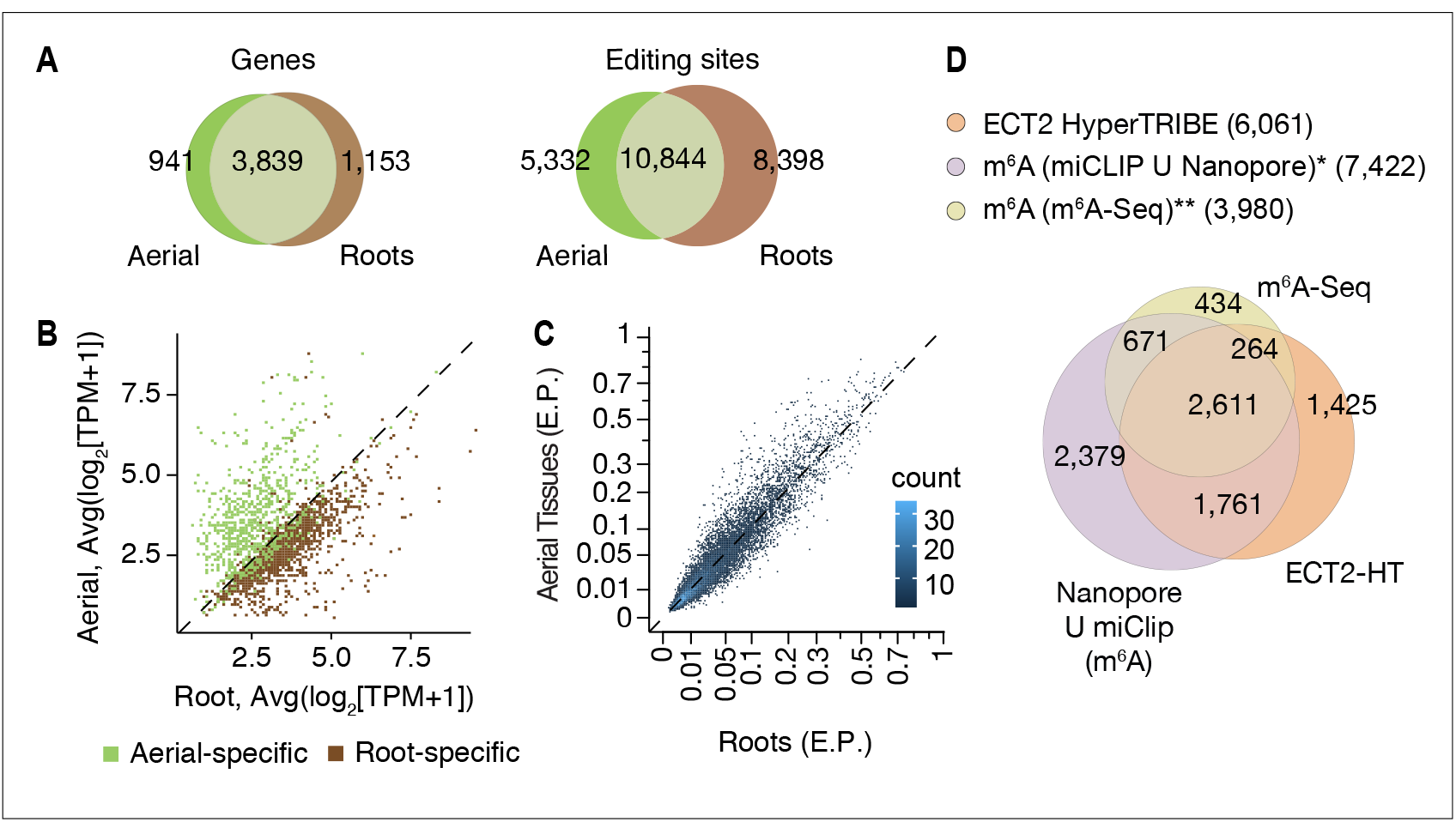
HyperTRIBE identifies m^6^A-reader targets in plants. **(A)** Overlap between ECT2-HT targets (genes and editing sites) in roots and aerial tissues, based on genes commonly expressed in both tissues. **(B)** Scatterplot showing the expression levels in roots and aerial tissues (mean log_2_(TPM+1) over the five ECT2-HT control samples) of the genes identified as aerial or root-specific targets. **(C)** Scatterplot of the editing proportions (E.P.) of significant editing sites in ECT2-HT for aerial vs root tissues. **(D)** Overlap between ECT2-HT targets and m^6^A-containing genes. *Parker et al. (2020); ** Shen et al. (2016).

### ECT2-mCherry can be specifically UV-crosslinked to target RNA in vivo

We next moved on to independent target and binding site identification via iCLIP (*Figure 3A*). We used transgenic lines expressing functional ECT2-mCherry under the control of the endogenous *ECT2* promoter in the *ect2-1* knockout background (Arribas-Hernández et al. 2018; Arribas-Hernández et al. 2020) to co-purify mRNAs crosslinked to ECT2 for iCLIP. Lines expressing the *ECT2^W464A^-mCherry* variant were used as negative controls, because this Trp-to-Ala mutation in the hydrophobic methyl-binding pocket of the YTH domain abrogates the increased affinity for m^6^A-RNA (Li et al. 2014b; Xu et al. 2014; Zhu et al. 2014). Accordingly, the point mutant behaves like a null allele in plants, despite its wild type-like expression pattern and level (Arribas-Hernández et al. 2018; Arribas-Hernández et al. 2020).

**Figure 3.**
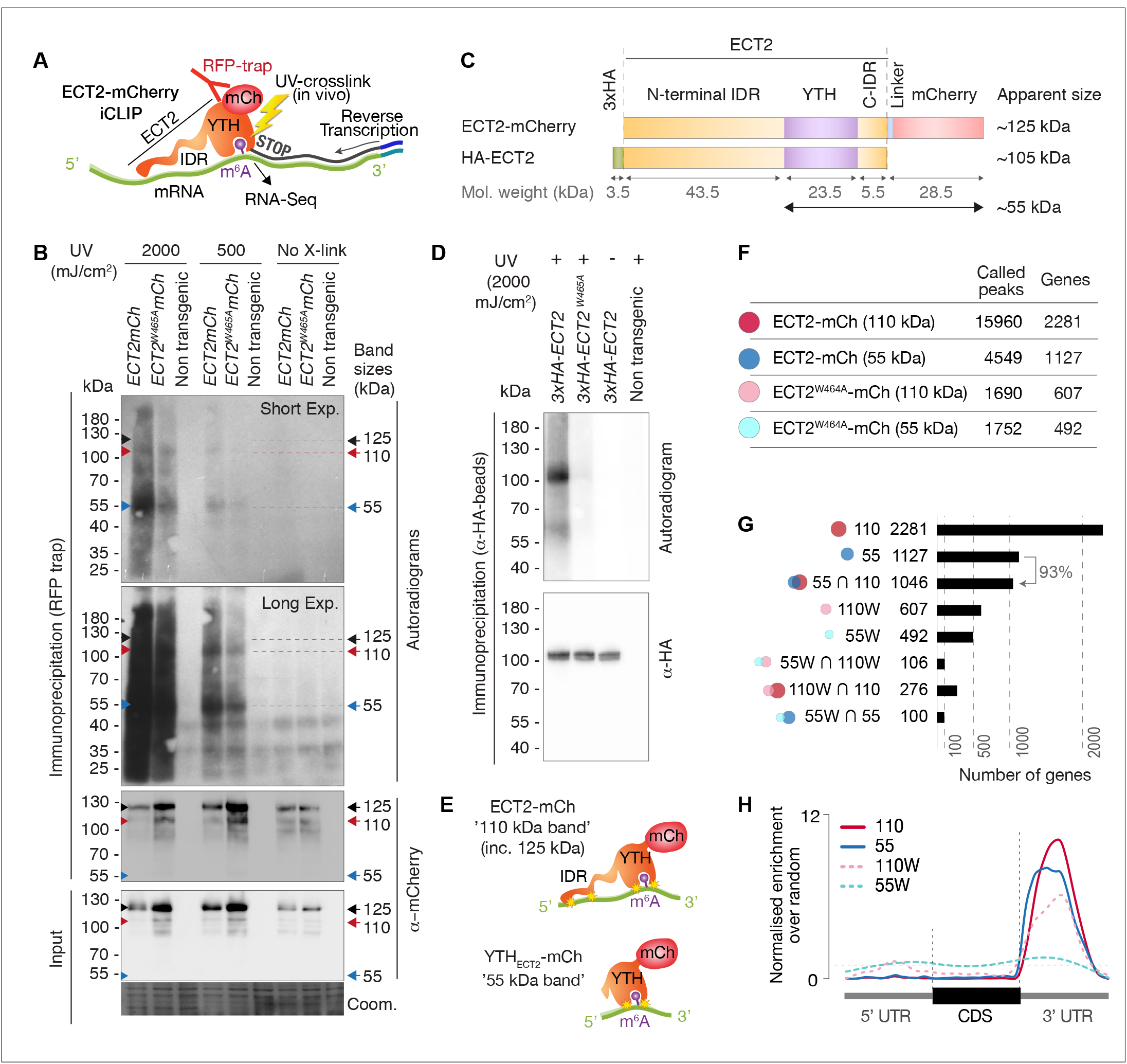
RNA-binding properties of ECT2 revealed by CLIP. **(A)** iCLIP experimental design. **(B)** Upper panels: autoradiogram (top) and α-mCherry protein blot (below) of RFP-trap immuno-purifications. Samples are cell extracts from 12-day-old seedlings expressing *ECT2-mCherry* or *ECT2^W464A^-mCherry* in the *ect2-1* mutant background after in vivo UV-crosslinking as indicated, and subjected to DNase digestion, partial RNase digestion, and 5’-^32^P labeling of RNA. Non-transgenic, Col-0 wild type. Lower panels: α-mCherry protein blot of the same extracts before immunopre-cipitation (input) and Coomassie staining of the membrane. Sizes corresponding to full length ECT2-mCherry (∼125 kDa) and the most apparent RNA bands are indicated with arrows. A repeat of the experiment with independently grown and crosslinked tissue is shown in the figure supplement 1A. **(C)** Schematic representation of ECT2-mCherry and HA-ECT2 fusion proteins with their apparent size (electrophoretic mobility). The molecular weight of each region is indicated. Notice that IDRs tend to show higher apparent sizes (low electrophoretic mobility) than globular domains. **(D)** Equivalent to B with lines expressing *3xHA-ECT2* variants in the *ect2-1* background, α-HA immuno-purifications and α-HA detection by western blot. **(E)** Cartoon illustrating the bands of labelled RNA co-purifying with ECT2-mCherry. Yellow stars indicate possible crosslinking sites. **(F)** Number of called peaks and genes detected from the 4 iCLIP libraries sequenced for this study (figure supplement 3). **(G)** Upset plot showing single and pairwise combinations of genes for the 4 sequenced iCLIP libraries. Additional intersections can be found in the figure supplement 4. **(H)** Metagene profiles depicting the enrichment along the gene body (5’UTR, CDS or 3’UTR) of the called iCLIP peaks detailed in F. **Figure supplement 1.** UV-crosslinked RNA co-purifies with ECT2-mCherry in a pattern that depends on the proteolytic cleavage of the ECT2 IDR. **Figure supplement 2.** Illustration of RNA-binding properties of ECT2 revealed by CLIP. **Figure supplement 3.** ECT2 iCLIP libraries. **Figure supplement 4.** Analysis of ECT2 iCLIP Libraries.

To test the feasibility of iCLIP, we first assessed the specificity of RNA co-purified with ECT2-mCherry after UV-illumination of whole seedlings by 5’-radiolabeling of the immunoprecipitated RNP complexes followed by SDS-PAGE. These tests showed that substantially more RNA co-purifies with wild type ECT2 than with ECT2^W464A^ upon UV-crosslinking, and that no RNA is detected without UV irradiation, or from irradiated plants of non-transgenic backgrounds (*Figure 3B, Figure 3—figure supplement 1A*). RNAse and DNAse treatments also established that the co-purified nucleic acid is RNA (*Figure 3—figure supplement 1B*). Thus, UV crosslinking of intact *Arabidopsis* seedlings followed by immunopurification successfully captures ECT2-RNA complexes that exist *in vivo*. Curiously, although the pattern of ECT2-RNA complexes with bands migrating at ∼110 and 55 kDa is highly reproducible, it does not correspond to the majority of the purified ECT2-mCherry protein which runs at ∼125 kDa in SDS-PAGE (*Figure 3B,C*). A variety of control experiments (*Figure 3—figure supplement 1C-E*), most importantly the disappearance of additional bands with use of an N-terminal rather than a C-terminal tag (*Figure 3C,D*), indicates that the band pattern arises as a consequence of proteolytic cleavage of the N-terminal IDR in the lysis buffer, such that fragments purified using the C-terminal mCherry tag include the YTH domain with portions of the IDR of variable lengths (*Figure 3—figure supplement 2*). Comparative analysis of RNA in 55 kDa and 110-125 kDa complexes may, therefore, provide insight into the possible role of the N-terminal IDR of ECT2 in mRNA binding (*Figure 3E*), an idea consistent with the comparatively low polynucleotide kinase labeling efficiency of full-length ECT2-mCherry-mRNA complexes (∼125 kDa) (*Figure 3B, Figure 3—figure supplement 2*). Thus, we prepared separate iCLIP libraries from RNA crosslinked to ECT2-mCherry/ECT2^W464A^-mCherry that migrates at ∼110-280 kDa (‘110-kDa band’), and at ∼55-75 kDa (’55-kDa band’) (*Figure 3—figure supplement 3*) to investigate the possible existence of IDR-dependent crosslink sites, and thereby gain deeper insights into the mode of YTHDF-binding to mRNA *in vivo*.

### ECT2-mCherry iCLIP peaks are enriched in the 3’-UTR of mRNAs

We identified a total of 15,960 iCLIP ‘peaks’ or crosslink sites (i.e. single nucleotide positions called by PureCLIP from mapped iCLIP reads (Krakau et al. 2017)) in 2,281 genes from the 110-kDa band of wild type ECT2-mCherry (henceforth referred to as ECT2 iCLIP peaks and targets, respectively). In the corresponding 55-kDa band, 4,549 crosslink sites in 1,127 genes were called, 93% of them contained in the 110-kDa target set (*Figure 3F,G, Figure 3—figure supplement 4, Supplementary file 2*). We note that these numbers perfectly agree with the idea of the 55-kDa band containing only YTH domain crosslink sites, while the full length may also include IDR crosslink sites. Importantly, for both libraries, the majority of crosslink sites mapped to the 3’-UTRs of mRNAs (*Figure 3H*, see *Figure 4A, Figure 4—figure supplement 1* for examples), coincident with the main location of m^6^A (*Figure 4B*) (Parker et al. 2020). Accordingly, the 3’-UTR specificity was largely lost in RNA isolated from 55-kDa ECT2^W464A^ (*Figure 3H*), for which neither YTH-domain nor IDR binding to RNA can be expected. Finally, iCLIP targets in full length (110-kDa band) ECT2 WT and ECT2^W464A^ overlapped only marginally (*Figure 3G*), providing molecular proof of the dependence of m^6^A-binding activity for ECT2 function. Nonetheless, the bias towards occurrence in the 3’-UTR was only reduced, not abolished, for crosslinks to the full-length ECT2^W464A^ protein, providing another indication that the IDR itself is able to associate with RNA-elements in 3’-UTRs (*Figure 3H*). We elaborate further on this important point by analysis of IDR-specific crosslinks to wild type ECT2 after in-depth validation of sets of ECT2 target mRNAs, and determination of the sequence motifs enriched around m^6^A and ECT2 crosslink sites.

**Figure 4.**
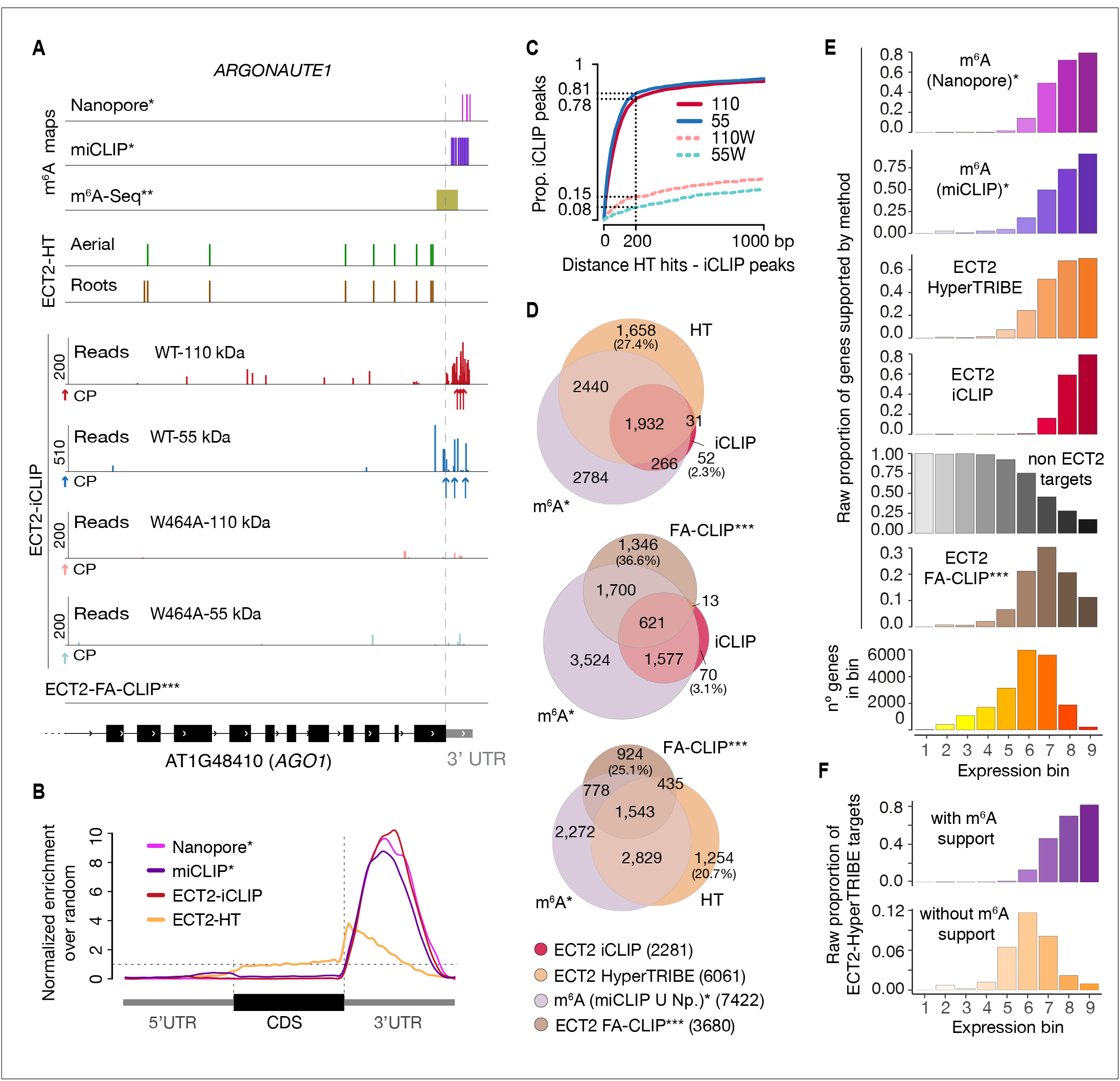
iCLIP identifies bona-fide ECT2 targets. **(A)** Example of an ECT2 target (*AGO1*) showing the distribution of m^6^A sites*, **, ECT2-iCLIP reads and peaks, ECT2-HT edited sites, and FA-CLIP peaks*** along the transcript. CP, called peaks. See more examples in the figure supplement 1. **(B)** Metagene profiles comparing the distributions along the gene body of ECT2-mCherry iCLIP peaks (wild type, 110-kDa band), ECT2-HT editing sites (in roots and aerial tissues) and m^6^A sites*. **(C)** Proportion of ECT2 iCLIP peaks within a given distance from the nearest ECT2-HT edited site. Numbers indicated on the y-axis show the proportion of ECT2 iCLIP peaks less than or equal to 200 nt from the nearest ECT2-HT edited site. **(D)** Overlap between genes supported as containing m^6^A or ECT2 targets by the different techniques indicated. The ECT2-HT target set includes the sum of targets identified in root and aerial tissues. Additional overlaps are shown in the figure supplement 2. **(E)** Proportions of genes in each expression bin either containing m^6^A or supported as ECT2 targets by the indicated techniques. **(F)** Proportion of ECT2-HT targets with or without support from m^6^A data (Nanopore*, miCLIP* or m^6^A-Seq**) in each expression bin. * Parker et al. (2020); ** Shen et al. (2016); *** Wei et al. (2018). **Figure supplement 1.** Distribution of m^6^A and ECT2 sites on ECT2 targets. **Figure supplement 2.** Overlaps between m^6^A-containing genes and ECT2-targets datasets. **Figure supplement 3.** Characteristics of ECT2-HyperTRIBE editing sites relative to target expression levels.

### iCLIP sites tend to be in the vicinity of HyperTRIBE editing sites

To evaluate the congruence of the results obtained by iCLIP and HyperTRIBE, we investigated the cumulative number of iCLIP sites as a function of distance to the nearest editing site determined by HyperTRIBE. This analysis showed a clear tendency for iCLIP peaks called with ECT2^WT^-mCherry, but not for ECT2^W464A^-mCherry, to be in the vicinity of editing sites (*Figure 4C*), indicating that the majority of called iCLIP peaks identify genuine ECT2 binding sites on mRNAs. Similar tendencies of proximity between iCLIP peaks and HyperTRIBE editing sites were previously observed for a *Drosophila* hnRNP protein (Xu et al. 2018). Although manual inspection of individual target genes confirmed these tendencies, it also revealed that ADAR-edited sites are too dispersed around iCLIP peaks to give precise information on the actual ECT2-binding sites (*Figure 4A, Figure 4—figure supplement 1*). Therefore, we used both HyperTRIBE and iCLIP for gene target identification, but relied on iCLIP peaks for motif analyses.

### ECT2 targets identified by iCLIP and HyperTRIBE overlap m^6^A–containing transcripts

To examine the quality of our target identification in further detail, we analyzed the overlap between ECT2 targets identified by iCLIP and HyperTRIBE. This analysis also included m^6^A mapping data obtained with either m^6^A-seq (Shen et al. 2016) or the single-nucleotide resolution methods miCLIP and Nanopore sequencing (Parker et al. 2020), as young seedlings were used in all cases. ECT2 targets identified by iCLIP and HyperTRIBE showed clear overlaps, both with each other and with m^6^A-containing transcripts, further supporting the robustness of ECT2 target identification via combined iCLIP and HyperTRIBE approaches (*Figure 4D upper panel, Figure 4—figure supplement 2*). Importantly, although some m^6^A-targets are expected not to be bound by ECT2 because of the presence of MTA in cells that do not express ECT2 (Arribas-Hernández et al. 2020), only 18% of the high-confident set of m^6^A-containing genes (with support from miCLIP and Nanopore) did not overlap with either ECT2 iCLIP or HT target sets (*Figure 4—figure supplement 2, arrow*). We also observed that HyperTRIBE identifies ∼3 times more ECT2 targets than iCLIP, possibly because of the bias towards high abundance inherent to purification-based methods like iCLIP (Wheeler et al. 2018). To test this idea, we compared the distribution of target mRNAs identified by the different techniques across 9 expression bins. As expected, a bias towards highly abundant transcripts was evident for iCLIP-identified targets compared to HyperTRIBE (*Figure 4E*). We also observed a similar bias for m^6^A-containing transcripts detected by miCLIP, another purification-based method, and in the Nanopore dataset (*Figure 4E*), probably explained by its relatively low sequencing depth (Parker et al. 2020). These observations also suggest that the higher sensitivity of HyperTRIBE (analyzed in detail in *Figure 4—figure supplement 3*) explains the lack of m^6^A support (by Nanopore or miCLIP) for 28% of ECT2 HT-targets (1,689) compared to only 4% (83) of ECT2 iCLIP targets (*Figure 4D, Figure 4—figure supplement 2, upper row*), since HT-targets may simply include genes that escape detection by m^6^A mapping methods due to low expression. Indeed, ECT2-HT targets without any m^6^A support were distributed in lower-expression bins compared to those with m^6^A support (*Figure 4F*). Intriguingly, ECT2 FA-CLIP targets (Wei et al. 2018) did not show a bias towards highly expressed genes, as their distribution over expression bins largely reflected that of the total number of genes (*Figure 4E*), and as many as 37% of FA-CLIP targets did not have m^6^A support (*Figure 4D, Figure 4—figure supplement 2, upper row*). In summary, these analyses show that ECT2 iCLIP and HT target sets are in excellent agreement with each other and with independently generated m^6^A maps, and that HyperTRIBE identifies targets below the detection limit of other techniques.

### ECT2 crosslink sites coincide with m^6^A miCLIP-sites and are immediately upstream of Nanopore m^6^A-sites

To characterize the sequence composition and exact positions of ECT2 binding sites relative to m^6^A, we first used the high resolution of iCLIP data to examine the position of ECT2 crosslink sites relative to m^6^A sites, determined at single-nucleotide resolution (Parker et al. 2020). This analysis showed that ECT2 crosslinks in the immediate vicinity, but preferentially upstream (∼11 nt) of Nanopore-determined m^6^A sites, with a mild depletion at the exact m^6^A site (*Figure 5A*, upper panel). Furthermore, while m^6^A-miCLIP sites corresponded to m^6^A Nanopore sites overall, a subset of m^6^A-miCLIP sites were located upstream of m^6^A-Nanopore sites and coincided well with ECT2-iCLIP peaks (*Figure 5A*). This pattern is probably explained by the fact that the UV illumination used in both iCLIP and miCLIP preferentially generates RNA-protein crosslinks involving uridine (Hafner et al. 2021), also detectable in the datasets analyzed here (*Figure 5B,C*). Thus, the depletion of ECT2-iCLIP sites at Nanopore-, but enrichment at miCLIP-determined m^6^A sites (*Figure 5A*) might be explained by the absence of uridine within the RR<u>A</u>C core of the m^6^A consensus motif, and perhaps also to some extent by reduced photoreactivity of the m^6^A base stacking with indole side chains of the YTH domain. Furthermore, the fact that nucleotides at -2, +1, and +2 positions are only expected to contribute sugar-phosphate backbone interactions with the YTH domain (Luo and Tong 2014; Theler et al. 2014; Xu et al. 2014), may also contribute to the absence of direct crosslinks at the m^6^A site relative to the adjacent bases.

**Figure 5.**
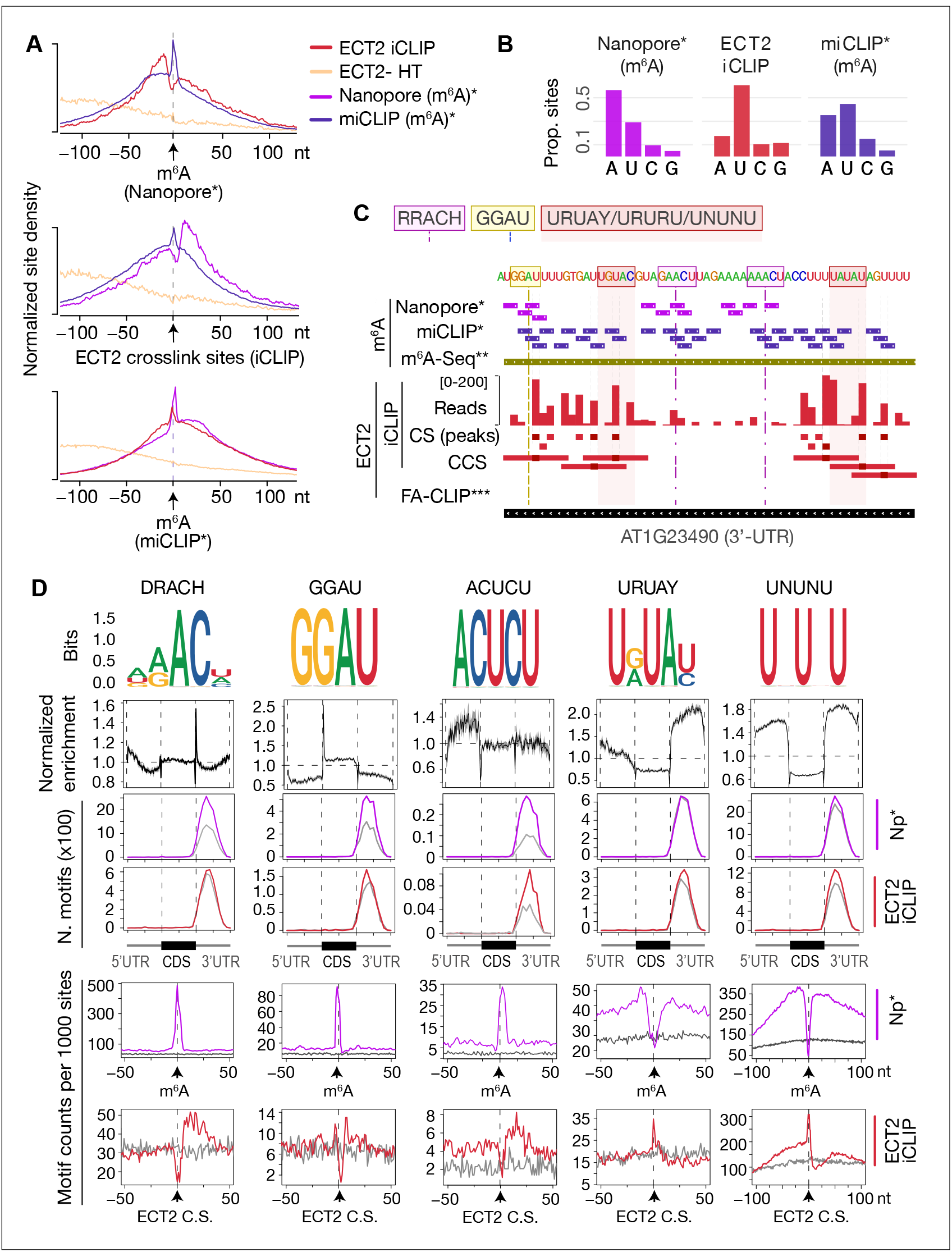
ECT2 UV-crosslinks to uridines in the immediate vicinity of DR(m^6^A)CH or GG(m^6^A)U sites. **(A)** Normalized density of sites at and up to +/-100 nt of either m^6^A Nanopore*, m^6^A miCLIP* or ECT2 iCLIP sites. **(B)** Proportion of m^6^A and ECT2 iCLIP sites at each nucleotide by the different methods. **(C)** View from IGV browser illustrating the presence of RRACH, GGAU and U-rich motifs in the vicinity of m^6^A and ECT2 sites in the 3’-UTR of AT1G23490 (*ARF1*). CS, crosslink sites; CSS, collapsed crosslink sites. **(D)** Key motifs analyzed in this study. From top to bottom: (1) motif logos for derived position weight matrices (PWMs); (2) normalized enrichment of motif locations across gene body; (3-4) total number of the relevant motif found at m^6^A-Nanopore* (3) or ECT2-iCLIP (4) sites according to gene body location. Grey lines indicate numbers found in a gene-body location-matched background set of sites of equivalent number; (5-6) distribution of the relevant motif relative to m^6^A-Nanopore* (5) or ECT2–iCLIP (6) sites. Grey lines represent the distribution for the same gene-body location-matched set as derived in the panels above. * Parker et al. (2020); ** Shen et al. (2016); *** Wei et al. (2018). **Figure supplement 1.** Sources of motifs and generation of position weight matrices. **Figure supplement 2.** Motif logos generated from position weight matrices **Figure supplement 3.** Enrichment of DRACH variants around m^6^A and ECT2 sites. **Figure supplement 4.** Uridines flanking DRACH result in additional motifs enriched at ECT2 iCLIP sites.

### DRACH, GGAU and U/Y-rich motifs are the most enriched around m^6^A/ECT2-sites

The 5’ shift observed for iCLIP and miCLIP sites relative to Nanopore sites might be explained by a higher occurrence of uridines upstream of m^6^A sites, a particularly interesting possibility given the numerous reports of U-rich motifs enriched around m^6^A sites in plants (Li et al. 2014c; Anderson et al. 2018; Miao et al. 2019; Zhang et al. 2019; Zhou et al. 2019; Luo et al. 2020) and animals (Patil et al. 2016). To investigate the sequence composition around m^6^A and ECT2 sites, we first performed exhaustive unbiased *de novo* motif searches using Homer (Heinz et al. 2010) (*Figure 5—figure supplement 1*) and extracted all candidate motifs, including the m^6^A consensus motif RRACH, as well as GGAU (Anderson et al. 2018), URUAY (Wei et al. 2018) and several other U-rich sequences. Combined with manually derived candidate motifs (*Figure 5—figure supplement 1B*), we then calculated position weight matrices (PWM) for a final set of 48 motifs and scanned for their occurrences genome-wide using FIMO (Grant et al. 2011) (*Figure 5—figure supplements 1,2*). This allowed us to determine three key properties. First, the global enrichment of the motifs at locations across the gene body. Second, the total count of occurrences of each motif at m^6^A sites and ECT2 iCLIP crosslink sites compared to a set of sites in non-target mRNAs matching the location within gene bodies of m^6^A/ECT2 iCLIP sites (expected background). Third, the distribution of the motifs relative to m^6^A and ECT2 iCLIP sites. The results of this systematic analysis (*Supplementary file 3*) were used to select those with a more prominent enrichment at or around m^6^A and ECT2 sites (*Figure 5D*). This approach defined two major categories of motifs of outstanding interest, RRACH-like and GGAU on the one side, and a variety of U/Y-rich motifs on the other. *Figure 5D* shows a minimal selection of such motifs, while a more comprehensive compilation is displayed in *Figure 5—figure supplements 3,4*. Not surprisingly, RRACH-like motifs were the most highly enriched at m^6^A sites and showed a clear enrichment immediately downstream of ECT2 crosslink sites in our analyses, with the degenerate variant DRACH being the most frequently observed (*Figure 5D, Figure 5—figure supplement 3*). Motifs containing GGAU behaved similarly to DRACH, with a sharp enrichment exactly at m^6^A sites and mild enrichment downstream of ECT2 peaks (*Figure 5D*), supporting a previous suggestion of GGAU as an alternative methylation site (Anderson et al. 2018). The possible roles of the U/Y-rich motifs in m^6^A deposition and ECT2 binding are analyzed in the following sections.

### Neighboring U/Ys results in enriched RRACH- and GGAU-derived motifs

We first noticed that several motifs retrieved around ECT2 crosslink sites by Homer constituted extended versions of **DRACH/GGAU** with *U*s upstream (e.g. *U***GAAC**/*U***GGAU**), or remnants of DR**ACH** with *U/C*s (*Y*s) downstream (e.g. **ACU***CU*). To test whether these motifs are indeed located adjacent to m^6^A, we examined their distribution and enrichment around ECT2 and m^6^A sites. The distributions showed a clear enrichment at m^6^A positions with a shift in the direction of the U/Y-extension (see *Figure 5D* for ACUCU and *Figure 5—figure supplement 4* for others). An enrichment over location-matched background sites close to ECT2-iCLIP sites was also apparent (see *Figure 5D* for ACUCU and *Figure 5—figure supplement 4* for others), further supporting that ECT2 preferentially crosslinks to uridines located in the immediate vicinity of DRACH (/GGAU). Thus, several enriched motifs around ECT2 crosslink sites are DRACH/GGAU-derived, and their detection in unbiased searches simply reflects a tendency of methylated DRACH/GGAU sites to be flanked by U/Y.

### Nature of U/Y-rich motifs more distant from m^6^A sites

U/R rich motifs without traces of adjacent DRACH (e.g. YUGUM, URUAY, URURU) showed a characteristic enrichment around, but depletion at, m^6^A sites. For some motifs, the enrichment was more pronounced 5’ than 3’ to m^6^A sites (see *Figure 5D* for URUAY and *Figure 5—figure supplement 4* for others). The distance between the site of maximal motif occurrence and the m^6^A site roughly coincided with the shift observed in ECT2 crosslink sites relative to m^6^A (*Figure 5A, upper panel*). Accordingly, these motifs were enriched exactly at ECT2 crosslink sites (see *Figure 5D* for URUAY and *Figure 5—figure supplement 4* for others), suggesting that they may constitute additional m^6^A-independent sites of interaction with ECT2. We also observed that the 3’ enrichment of YYYYY was closer to m^6^A than that of UUUUU/URURU/URUAY (*Figure 5—figure supplement 4, 2^nd^ row from the top*), indicating a preference for hetero-oligopyrimidine tracts immediately downstream the m^6^A site, as suggested by the 3’-enrichment of DRACUCU-type motifs as described above.

Taken together, these results suggest that *N6*-adenosine methylation preferentially occurs in DRACH/GGAU sequences surrounded by stretches of pyrimidines, with a preference for YYYYY (e.g. CUCU) immediately downstream, URURU (including URUAY) immediately upstream, and UUUUU/UNUNU slightly further away in both directions. The enrichment of ECT2 crosslink sites at these motifs, and the fact that the m^6^A-binding deficient mutant of ECT2 (W464A) crosslinks preferentially to 3’UTRs through its N-terminal IDR, indicate IDR-mediated binding to U/R- and Y-rich motifs around m^6^A.

### DRACH/GGAU motifs are determinants of m^6^A deposition at the site, while flanking U(/Y)-rich motifs are indicative of m^6^A presence and ECT2 binding

Since our analysis thus far uncovered several motifs of potential importance for m^6^A deposition and ECT2 binding, we employed machine learning to distinguish m^6^A and ECT2 iCLIP sites from random location-matched background sites using motif-based features. Importantly, the underlying classification model includes all motif features within the same model, allowing an evaluation of the importance of the motifs relative to each other. We used as features the number of matches to each of the 48 motifs (*Figure 5—figure supplement 2*) in three distinct regions relative to the methylated site according to Nanopore sequencing (Parker et al. 2020), defined as position 0: “at” [-10 nt; +10 nt], “down” [-50 nt; -10 nt], or “up” [+10 nt; +50 nt] (*Figure 6A*). The model involving all motifs could successfully distinguish the methylated sites from the background as indicated by an area under the receiver operating characteristic curve (true positive rate versus false positive rate, AUC) of 0.93, and even a reduced model incorporating only the top 10 features from the full model classified sites largely correctly (AUC = 0.86; *Figure 6—figure supplement 1*). The top 16 features ordered by importance from the full model confirmed that RRAC/DRACH or GGAU at the site was indicative of the presence of m^6^A (*Figure 6B*). Interestingly, U/Y-rich sequences (UNUNU and YYYYY in particular) flanking the site were also strongly indicative (*Figure 6B*). Some motifs showed a skew in their feature importance score, with UNUNU and YUGUM showing a preference to be upstream, and YYYYY downstream (*Figure 6B*), thus corroborating our previous observations (*Figure 6C*).

**Figure 6.**
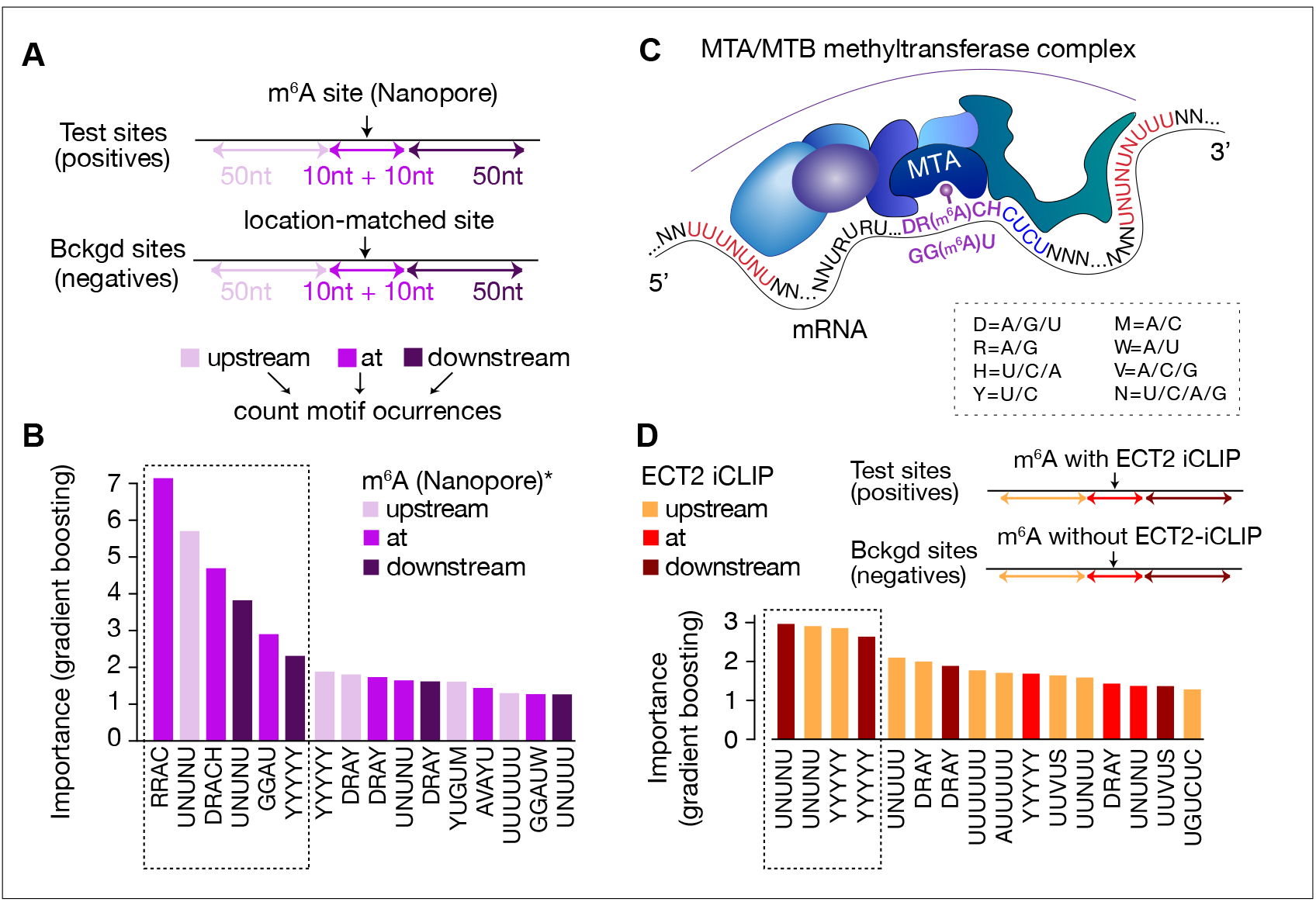
Distal U-rich motifs and at-the-site DRACH/GGAU are determinants for m^6^A deposition. **(A)** Diagram representing strategy for machine learning model trained to distinguish m^6^A Nanopore* sites from their respective gene-body location matched background sets. **(B)** Bar plots showing top 16 motif feature importance scores from the m^6^A model, ordered from left to right by importance. Dotted rectangle highlights motifs with outstanding importance compared to the rest. **(C)** Cartoon representing the most important motifs found at and around m^6^A sites. UPAC-IUB codes to define multiple nucleotide possibilites in one position are indicated. **(D)** Machine learning model trained to distinguish between m^6^A sites with and without ECT2 crosslink sites, and the resulting bar plot showing top 16 motif feature importance scores. Nucleotide distances for intervals, order and dotted box are as in A. * Parker et al. (2020). **Figure supplement 1.** Model performance ROC curves for distinguishing sequence preferences of m^6^A or ECT2-bound sites.

We used a similar modeling approach to identify non-m^6^A determinants of ECT2 binding, in this case comparing m^6^A sites within 10nt-distance of ECT2-iCLIP sites to m^6^A sites without ECT2-iCLIP sites nearby (AUC=0.94, and AUC=0.84 using only the top 10 features, *Figure 6—figure supplement 1*). In agreement with previous observations, this model showed flanking U/Y-rich sequences as the main determinants for ECT2 crosslinking (*Figure 6D*).

### The U(-R) paradox: URURU-like sequences around m^6^A sites repel ECT2 binding, while U-rich sequences upstream enhance its crosslinking

To investigate the idea of URURU-like motifs as additional sites of ECT2 binding upstream of the m^6^A-YTH interaction site, we split Nanopore-m^6^A sites according to two criteria: 1) whether they occur in ECT2-target transcripts (both permissive and stringent sets analyzed separately), and 2) for ECT2 targets, whether there is an ECT2 crosslink site within 25 nt of the m^6^A site (‘near’) or not (‘far’). Although there was no obvious difference between these categories for most of the motifs (*Supplementary file 3, page 2*), some U-rich sequences displayed distinctive features (*Figure 7A, Figure 7—figure supplement 1*) that can be summarized as follows. If a transcript has m^6^A <u>and</u> ECT2 sites in close proximity, it is: i) more likely to have UNUNU/UUUUU/YYYYY sequences upstream of the m^6^A site than targets with distantly located ECT2 binding sites or than non-ECT2 targets; ii) less likely to have UUUUU/URURU sequences downstream of the m^6^A site, possibly because ECT2 prefers CUCU-like sequences downstream; iii) less likely to have URURU/URUAY-like motifs upstream of the m^6^A site. The latter observation is striking, because for the specific subset of ECT2-bound m^6^A-sites with URURU/URUAY upstream of m^6^A, these sequences tend to crosslink to ECT2, as seen by the enrichment spike at ECT2 crosslink sites (*Figure 5D, Figure 7— figure supplement 1, bottom panels*). Although these two results seem contradictory at first glance, they may be reconciled by a model in which a URURU/URUAY-binding protein would compete with ECT2 for binding adjacent to m^6^A. If that protein is absent, ECT2 may bind to the site, potentially via its IDR, to stabilize the low-affinity YTH-m^6^A interaction and crosslink efficiently due to the U-content. Conversely, if occupied by the alternative interacting protein, the site might repel ECT2 (see discussion and *Figure 7B*).

**Figure 7.**
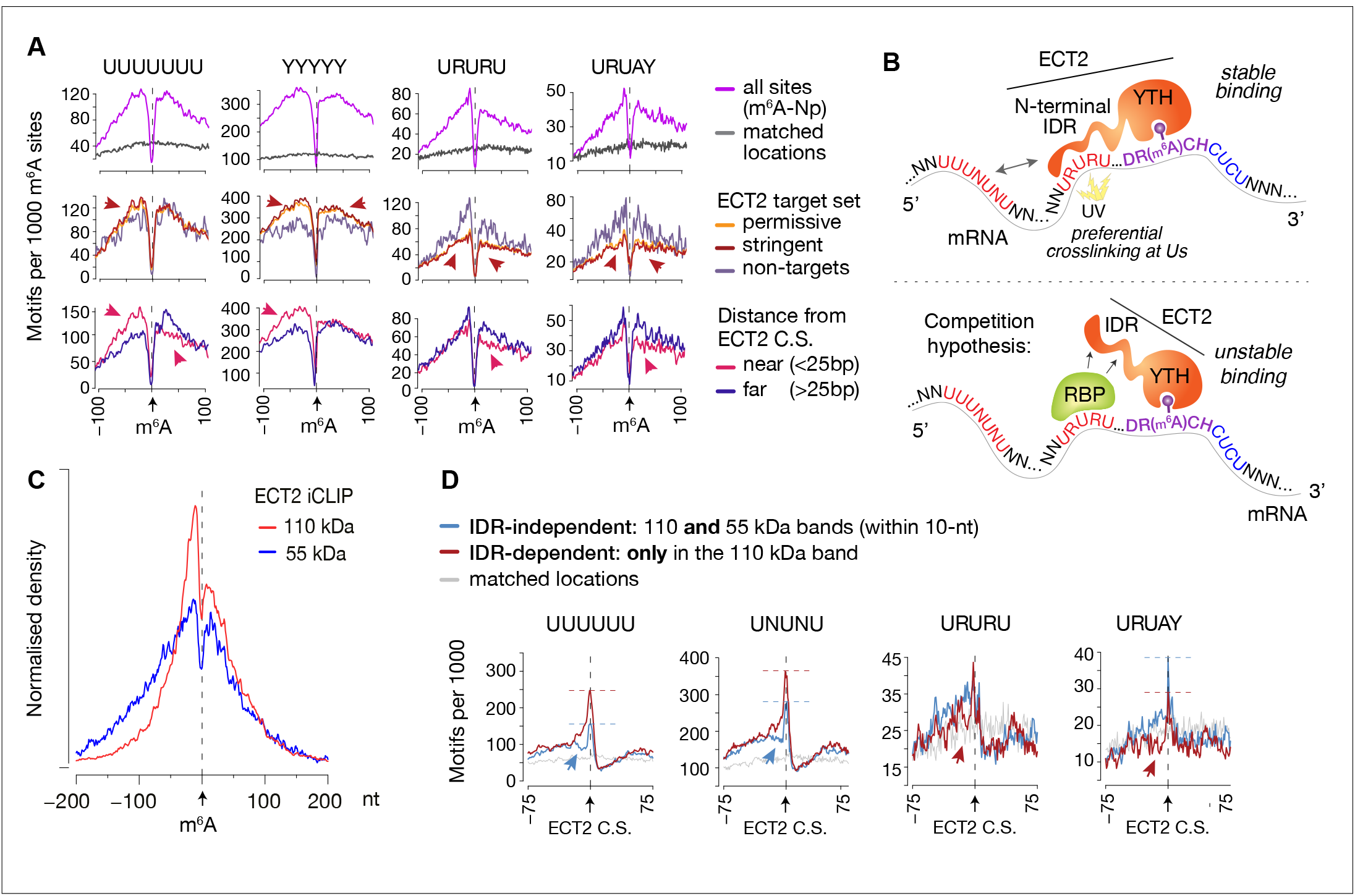
IDR-dependent binding of ECT2 to U-rich motifs 5’ of m^6^A. (**A**) Top panels: Distance-based enrichment of motifs at and around m^6^A-Nanopore (Np, Parker et al., 2020) sites, plotted as motif counts per 1000 m^6^A sites (purple lines). Grey lines indicate the enrichment in a location-matched background set as in Figure 5D. Middle and bottom panels: sites are split according to whether they sit on ECT2 targets (middle), or to distance from the nearest ECT2 crosslink site (for ECT2-iCLIP targets only) (bottom). Additional motifs are shown in the figure supplement 1. (**B**) Cartoon illustrating the ECT2 IDR RNA-binding and competition hypotheses. (**C)** Normalized density of ECT2 iCLIP crosslink sites identified in the libraries corresponding to the 110 and 55 kDa bands (Figure 3B) at and up to +/-200 nt of m^6^A Nanopore* sites. (**D**) Motifs per 1000 ECT2 iCLIP crosslink sites (CS) split according to whether they are found in libraries from both 110 kDa and 55 kDa bands (IDR-independent’), or exclusively (distance > 10 nt) in the 110 kDa band (’IDR-dependent’). Grey lines indicate the enrichment in a location-matched background set as in Figure 5D. Additional motifs are shown in the figure supplement 2 and supplementary file 3. **Figure supplement 1.** Motif preferences around m^6^A sites according to ECT2 binding. **Figure supplement 2.** Dependency of the ECT2 IDR for motif enrichment.

### The N-terminal IDR of ECT2 is involved in preferential crosslinking at U-rich sequences and URURU-repulsion immediately upstream m^6^A sites

We reasoned that insights into contacts between ECT2 and mRNA may be gained by analysis of the iCLIP libraries prepared with the ‘YTH-mCherry’ truncation devoid of the N-terminal IDR (‘55 kDa band’) compared to the full-length ECT2-mCherry (‘110 kDa’) (*Figure 3, Figure 3—figure supplements 2-4*). Initial inspection of the distribution of ECT2 peaks relative to Nanopore-m^6^A sites showed that the 5’-3’ asymmetry observed with full-length ECT2 was largely reduced with the truncated protein (*Figure 7C*), as was the bias towards uridines (*Figure 7—figure supplement 2A*). These observations suggest that the IDR indeed is implicated in binding to U-rich regions upstream of m^6^A. We next split the full-length ECT2 iCLIP peaks according to whether they are present in libraries from both full length and truncated forms (’IDR-independent’), or exclusively in the full length (’IDR-dependent’) (distance > 10 nt), and plotted the enrichment of the studied motifs relative to the crosslink site (*Figure 7D, Figure 7—figure supplement 2B; Supplementary file 3-page 2*). UUUUUU/UNUNU-like motifs were more enriched at and immediately upstream of IDR-dependent crosslink sites relative to the IDR-independent ones, supporting preferential crosslinking of the IDR to Us in this region. Remarkably, the exact opposite was true for URURU/URUAY motifs that showed modest depletion 5’ to IDR-dependent crosslink sites relative to their IDR-independent counterparts (*Figure 7D*). These observations are consistent with a model of an RNA binding protein competing with the ECT2 IDR for interaction with upstream URURU/URUAY motifs (*Figure 7B*).

## Discussion

### Methodology for mapping protein-RNA interactions in plants

Our work establishes experimental and computational approaches to implement HyperTRIBE for unbiased and sensitive mapping of direct targets of RNA binding proteins in plants. Two points are particularly relevant in this regard. First, the examples studied here show that stable transgenic expression of *Dm*ADARcd does not lead to detrimental phenotypes, perhaps because of the generally low editing proportions obtained *in vivo*. Second, the rigorous statistical approach developed to call editing sites makes HyperTRIBE powerful, despite the low editing proportions observed. We also note that ECT2 is well suited to verify that HyperTRIBE mostly recovers directly bound target RNAs, because of the possibility to cross-reference the data with independently obtained m^6^A maps (Parker et al. 2020). The combination of iCLIP and HyperTRIBE for unbiased mapping of targets proved particularly attractive for at least two reasons. First, the convergence on overlapping target sets by orthogonal methods strengthens the confidence that the identified targets are biologically meaningful. Second, HyperTRIBE, especially with the novel computational approach for calling of editing sites developed here, offers higher sensitivity than iCLIP, while iCLIP is unmatched in providing information on binding sites within target RNAs. It is possible that better positional information on binding sites may be obtained from HyperTRIBE data using maximal editing proportions rather than statistical significance as the parameter to call editing sites. Indeed, recent work on the use of HyperTRIBE to identify targets of the RNA-binding protein MUSASHI-2 (MSI-2) in leukemic stem cells recovered the known MSI-2 binding site as enriched around editing sites in targets (Nguyen et al. 2020). Nonetheless, our data shows that highly edited sites match the ADAR substrate consensus site better than lowly edited sites, suggesting that site proximity to ADAR is not the only determinant of editing proportions. Finally, our work also clearly indicates that FA-CLIP, now used in at least two studies involving YTH domain proteins (Wei et al. 2018; Song et al. 2021), is not a recommendable technique, as it recovers many false positives and fails to include many genuine targets. Thus, with the possible exception of cases in which evidence for indirect association is specifically in demand, such as the recent study in human cells of mixed tailing of viral RNA by the cellular terminal nucleotidyltransferase TENT4 (Kim et al. 2020), FA-CLIP should not be used for identification of RNAs associating with a particular RNA-binding protein of interest.

### Core elements in m^6^A writing: DRACH, GGAU and U/Y-rich motifs

Our analyses of motif enrichments around m^6^A and ECT2 crosslink sites clarify roles of previously reported motifs and uncovers new motifs of importance in m^6^A writing and ECT2 binding. Since m^6^A is a prerequisite for ECT2 binding, any analysis of determinants of ECT2 binding must consider determinants of *N6*-adenosine methylation separately. Three conclusions stand out from our analysis in this regard. First, the major *N6*-adenosine methylation site is DRACH, consistent with conclusions from multiple other studies. Second, GGAU is a minor *N6*-adenosine methylation site, as seen by its enrichment directly at m^6^A-sites. Third, m^6^A occurs in DRACH/GGAU islands embedded in U-rich regions. Such U-rich regions around m^6^A sites emerged from sorting of methylated from non-methylated transcripts by machine learning as being of similar importance for recognition of m^6^A-containing transcripts from sequence features as DRACH and GGAU at m^6^A sites, suggesting their implication in MTA/MTB-catalyzed adenosine methylation (*Figure 6C*). This, in turn, may also explain the pronounced 3’-UTR bias of m^6^A occurrence, as extensive poly-U and poly-pyrimidine tracts are rare in coding regions (*Figure 5D, 2^nd^ row; Supplementary file 3-page 1)*. As a special case in this context, our analyses suggest a simple explanation for the tendency of m^6^A to occur at stop codons. UAA and UGA correspond to DRA, increasing the frequency of occurrence of DRACH directly at stop codons (*Figure 5D, 2^nd^ row)*, many of which have adjacent U-rich elements in the 3’-UTRs. We note that the observed pattern is in agreement with a role of the poly(U)-interacting proteins RBM15A/B associated with the methyltransferase complex in mammalian cells in guiding methylation (Patil et al. 2016). Whether a similar mechanism operates in plants, potentially via the distant RBM15A/B homologue FPA (Arribas-Hernández and Brodersen 2020), remains to be investigated.

### Reading of DR(m^6^A)CH in 3’-UTRs of target mRNAs by ECT2

It is a major conclusion of the present work that ECT2 binds to m^6^A predominantly in the DR(m^6^A)CH sequence context *in vivo*, consistent with reading of m^6^A written by the conserved nuclear MTA/MTB methyltransferase. This key conclusion refutes the claim by Wei et al. (2018) that ECT2 binds to the supposedly plant-specific m^6^A-containing sequence motif URU(m^6^A)Y, and it thereby reconciles knowledge on m^6^A-YTHDF axes in plants specifically and in eukaryotes more broadly. The phenotypic similarity of plants defective in MTA/MTB writer and ECT2/3/4 reader function is now coherent with the locations of MTA/MTB-written m^6^A and ECT2 binding sites transcriptome-wide, and it is now clear that plants do not constitute an exception to the general biochemical framework for eukaryotic m^6^A-YTHDF function in which YTHDF proteins read the m^6^A signal written by the MTA/MTB methyltransferase.

### The role of U-rich motifs 5’ to m^6^A sites in ECT2 binding: direct interaction of the IDR of ECT2 with mRNA

The pronounced protease-sensitivity of IDRs, leading to limited proteolysis of ECT2 upon cell lysis after *in vivo* crosslinking allowed us to extract information on the mode of ECT2-RNA binding from different observations, all converging on the conclusion that the IDR of ECT2 participates in RNA binding. First, RNA-complexes with YTH-mCherry were 5’-labeled by polynucleotide kinase much more efficiently than RNA-complexes with full-length ECT2-mCherry, indicating that the IDR limits accessibility to the 5’ of bound mRNAs. Second, in contrast to the m^6^A-binding deficient YTH^W464A^-mCherry truncation, the full-length ECT2^W464A^-mCherry mutant retained an enrichment of crosslink sites in 3’-UTRs. Third, crosslinks specific to the IDR (i.e. observed only with full-length ECT2-mCherry-RNA complexes, but not with YTH-Cherry-RNA complexes) could be assigned, and have two notable properties. They are mainly 5’ to m^6^A sites, and thereby cause a conspicuously asymmetric distribution of ECT2 crosslink sites around m^6^A sites, not seen with crosslinks to the YTH-mCherry fragment. In addition, the IDR-specific crosslinks are specifically enriched in U-rich elements of the type UUUUUU and UNUNU immediately upstream. Taken together, these observations suggest that the IDR of ECT2 participates in locating ECT2 to 3’-UTRs by association with U-rich elements. Thus, ECT2, and perhaps YTHDF proteins more generally given their highly similar YTH domains, appears to bind RNA through multivalent interactions among which the YTH domain is responsible for m^6^A-binding, and the IDR is responsible for interaction with adjacent elements. We note that the notion of RNA-interaction by IDRs has precedent (Corley et al. 2020), is consistent with the modest affinity of isolated YTHDF domains for m^6^A-containing oligonucleotides (Patil et al. 2018), and is reminiscent of the recent demonstration that transcription factors use their globular DNA-binding domains to recognize core sequence elements of promoters, and their IDRs to provide additional DNA contacts, contributing to specificity (Brodsky et al. 2020). We stress that although our data point to an important role of the IDR in RNA binding, it does not in any way suggest that this is the only function of the IDR, and protein-protein interactions involving the IDR are likely to be key to understanding YTHDF function molecularly.

### URUAY as sites of competitive interaction between ECT2 and other RNA-binding proteins

Despite the conclusions that URUAY does not contain m^6^A in Arabidopsis, and that ECT2 binds to DR(m^6^A)CH, our detailed analysis of sequence motifs enriched around m^6^A and ECT2 iCLIP crosslink sites shows that additional motifs, including URUAY, are likely to be implicated in m^6^A reading by ECT2, even if not directly. In contrast to other m^6^A-proximal, pyrimidine-rich sequences (e.g. UNUNU, YYYYY) that may be of importance for both m^6^A writing and ECT2 binding, URUAY appears to have ties more specifically to ECT2 binding thanks to three properties. (1) When present 5’ to m^6^A sites, it crosslinks to ECT2, suggesting that some part of the protein can be in contact with URUAY. (2) URUAY is more enriched close to m^6^A-sites for which there is no evidence of ECT2-binding, suggesting that it weakens ECT2 binding. This latter point is also consistent with the distinction of ECT2-bound from non-ECT2 bound m^6^A sites by machine learning that did not find URUAY to be of importance for ECT2-bound sites. (3) The URUAY enrichment 5’ to ECT2 crosslink sites is observed only when crosslinks to both full-length protein and the YTH-mCherry fragment are considered (IDR-independent), but disappears when crosslinks specific to the full-length protein (IDR-dependent) are analyzed. Although these observations may be explained by multiple scenarios, we find a simple, yet at present speculative, model attractive: URUAY may be a site of competition between the IDR of ECT2 and another, as yet unknown, RNA binding protein. We also note that the idea of a URUAY-binding protein influencing ECT2-binding and/or regulation is consistent with the recovery of formaldehyde crosslinks between ECT2 and URUAY (Wei et al. 2018), in this case indirectly. Finally, it is intriguing that URUAY resembles part of a Pumilio binding site (Hafner et al. 2010; Huh et al. 2013), raising the tantalizing possibility of functional interaction between YTHDF and Pumilio proteins. In any event, the functional dissection of the URUAY element in m^6^A-reading now constitutes a subject of major importance, emphasized by the broad conservation of its enrichment around m^6^A sites across multiple plant species, including rice (Li et al. 2014c; Zhang et al. 2019), maize (Luo et al. 2019; Miao et al. 2019), tomato (Zhou et al. 2019), and Arabidopsis (Miao et al. 2019).

## Methods

All data analyses were carried out using TAIR 10 as the reference genome and Araport11 as the reference transcriptome. Unless otherwise stated, data analyses were performed in R (https://www.R-project.org/) and plots generated using either base R or ggplot2. (https://ggplot2.tidyverse.org).

### Plant material

All lines used in this study are in the *Arabidopsis thaliana* Col-0 ecotype. The mutant alleles or their combinations: *ect2-1* (SALK_002225) (Arribas-Hernández et al. 2018; Scutenaire et al. 2018; Wei et al. 2018)*, ect3-1* (SALKseq_63401)*, ect4-2* (GK_241H02), and *ect2-1/ect3-1/ect4-2* (*te234*) (Arribas-Hernández et al. 2018) have been previously described. The transgenic lines *ECT2pro:ECT2-mCherry-ECT2ter*, *ECT2pro:ECT2^W464A^-mCherry-ECT2ter, ECT2pro:3xHA-ECT2-ECT2ter*, or *ECT2pro:3xHA-ECT2^W464A^-ECT2ter* have also been described or generated by floral dip in additional mutant backgrounds using the same plasmids and methodology (Arribas-Hernández et al. 2018; Arribas-Hernández et al. 2020).

### Growth conditions

Seeds were surface-sterilized, germinated and grown on vertically disposed plates with Murashige and Skoog (MS)-agar medium (4.4 g/L MS, 10 g/L sucrose, 10 g/L agar) pH 5.7 at 20°C, receiving ∼70 μmol m^-2^ s^-1^ of light in a 16 hr light/8 hr dark cycle as default. To assess phenotypes of adult plants, ∼8-day-old seedlings were transferred to soil and maintained in Percival incubators also under long day conditions. Additional details and variations of growth conditions for specific experiments can be found in Supplemental Methods.

### Generation of transgenic lines for HyperTRIBE

We employed USER cloning (Bitinaite and Nichols 2009) to generate *ECT2pro:ECT2-FLAG-ADAR-ECT2ter* and *ECT2pro:FLAG-ADAR-ECT2ter,* constructs in pCAMBIA3300U (pCAMBIA3300 with a double PacI USER cassette inserted between the *Pst*I-*Xma*I sites at the multiple cloning site (Nour-Eldin et al. 2006)). Details on the cloning procedure can be found in Supplemental Methods. *Arabidopsis* stable transgenic lines were generated by floral dip transformation (Clough and Bent 1998) of Col-0 WT, *ect2-1*, or *te234*, and selection of primary transformants (T1) was done on MS-agar plates supplemented with glufosinate ammonium (Fluka) (10 mg/L). We selected 5 independent lines of each type based on segregation studies (to isolate single T-DNA insertions), phenotypic complementation (in the *te234* background) and transgene expression levels assessed by FLAG western blot (see Supplemental Methods).

### HyperTRIBE library preparation

We extracted total RNA (see Supplemental Methods) from manually dissected root tips and apices (removing cotyledons) of 5 independent lines (10-day-old T2 seedlings) of each of the lines used for ECT2-HT, to use as biological replicates. Illumina mRNA-Seq libraries were then prepared by Novogene (see Supplemental Methods) after mRNA enrichment with oligo(dT) beads (18-mers).

### HyperTRIBE editing site calling

Significant differentially edited sites between *ECT2-FLAG-ADAR* (fusion) and *FLAG-ADAR* (control) samples were called according to our hyperTRIBE**R** pipeline (https://github.com/sarah-ku/hyperTRIBER), testing all nt positions with some evidence of differential editing across multiple samples. Significant (adjusted-*p*-value < 0.01 and log_2_FC > 1) A-to-G hits were further filtered and annotated according to Araport11 by integrating quantification information generated using Salmon (Patro et al. 2017), based on the Araport11 transcriptome with addition of *FLAG-ADAR* sequence. See Supplemental Methods for additional details.

### Analysis of HyperTRIBE sites

For all significant editing sites (sig. E.S.), editing proportions (E.P.) where calculated as G/(A+G) where A, G are the number of reads covering the E.S. with A or G at the site, respectively. Sample-specific E.P.s for all sig. E.S. were used for principal component analyses and correlations between *FLAG-ADAR* TPM and E.P., and replicate-averaged E.P. was used for density plots, condition- or cell-type-based comparisons and comparisons over expression bins. For expression bins, the log_2_(TPM+1) values for all expressed genes in either aerial tissues, roots or combined were split into 9 bins of increasing expression. The Support of ECT2 target or m^6^A gene sets was calculated by the proportion of genes in a given bin out of the total number of genes in that bin. See Supplemental Methods for additional details and methods.

### CLIP experiments and iCLIP library preparation

*In vivo* UV-crosslinking of 12-day-old seedlings and construction of iCLIP libraries were optimized for ECT2-mCherry from the method developed by Prof. Staiger’s group for Arabidopsis GRP7-GFP (Meyer et al. 2017; Köster and Staiger 2020). Details can be found in Supplemental Methods.

### iCLIP data analysis and peak calling

Sequenced reads were mapped to TAIR10 after being processed by trimming, demultiplexing and discarding short reads. Peak calling of uniquely mapped reads was done using PureCLIP 1.0.4 (Krakau et al., 2017) after removal of PCR duplicates. Gene annotation of peaks was carried out using the hyperTRIBE**R** pipeline. Details can be found in Supplemental Methods.

### Analysis of publicly available data

Single cell expression data and marker genes associated with 15 clusters annotated to cell types in roots was downloaded from Denyer et al. (2019). Single nucleotide resolution locations of m^6^A sites (defined according to Nanopore or miCLIP) were downloaded from Parker et al., 2020. Intervals defining m^6^A locations based on m^6^A-seq were downloaded from Shen et al. 2016, and intervals defining locations of ECT2-bound sites as determined by FA-CLIP were downloaded from Wei et al., 2018. For consistency with HyperTRIBE and ECT2-iCLIP, all sets of m^6^A or ECT2-bound sites were gene annotated using the hyperTRIBE**R** pipeline.

### Motif analysis

A list of 48 motifs was compiled from multiple sources and for each motif a position weight matrix (PWM) was generated based on local sequence frequencies around ECT2-iCLIP peaks and used as input to FIMO (Grant et al. 2011) to detect genome-wide occurrences. In order to account for location-specific sequence contexts (typically 3’UTR), each site from iCLIP or m^6^A (Parker et al. 2020) sets was assigned a random ‘matched background’ site, in a non-target gene, at the same relative location along the annotated genomic feature (5’UTR, CDS or 3’UTR) of the site. Distributions of motifs per 1000 sites over distance, centering on ECT2-iCLIP or m^6^A sites and the respective matched backgrounds were generated using a custom R-script (https://github.com/sarah-ku/targets_arabidopsis). Sets were further split according to IDR or target status (see Supplemental Methods for further details).

For machine learning, features were generated from motifs according to their relative locations in windows from m^6^A or ECT-iCLIP sites. Importance scores were generated using gradient boosting *gbm* (https://github.com/gbm-developers/gbm), with performance statistics based on the AUC calculated from held-out data. See Supplemental Methods for additional details.

## Data Access

### Accession numbers

The raw and processed data for HyperTRIBE (ECT2-HT) and ECT2-iCLIP have been deposited in the European Nucleotide Archive (ENA) at EMBL-EBI under the accession number PRJEB44359.

### Code availability

The code for running the hyperTRIBE**R** pipeline and for bioinformatics analyses is available at https://github.com/sarah-ku/targets_arabidopsis.

## Acknowledgements

We thank Lena Bjørn Johansson and Phillip Andersen for technical assistance in the construction of transgenic lines, and Theo Bølsterli, René Hvidberg Petersen and their teams for plant care. Kim Rewitz is thanked for providing the *Drosophila* larvae and flies used for cDNA extraction to clone *Dm*ADARcd. We acknowledge Maria Louisa Vigh for cloning of FLAG-*Dm*ADARcd, Katja Meyer and Kristina Neudorf for support during iCLIP library construction in Bielefeld, and Simon Bressendorff and Mathias Tankmar for experimental support. This work was supported by a Consolidator Grant from the European Research Council (PATHORISC, ERC-2016-COG 726417) and a Research Grant from the Independent Research Fund Denmark (9040-00409B) to P.B.; an EMBO Short Term Fellowship (STF 7614) to L.A.-H.; a Research Grant from DFG (STA653/14-1) to D.S.; and a Starting Grant from the European Research Council (638173) and a Sapere Aude Starting Grant from the Independent Research Fund Denmark (6108-00038B) to R.A.

## Author Contributions

P.B. and L.A.-H. designed and coordinated the study. L.A.-H. built the biological material for HyperTRIBE and S.R. developed the hyperTRIBE**R** pipeline, called edited sites to define HyperTRIBE target sets and assessed their veracity. L.A.-H. and T.K. performed iCLIP experiments and produced iCLIP libraries, M.L. analyzed iCLIP data, C.P. and S.R. executed *de novo* motif discovery. and S.R. studied the overlap between m^6^A and ECT2 target sets and performed motif enrichment analyses. R.A. supervised work related to hyperTRIBE**R** development and sequence motif analysis and D.S. supervised work related to iCLIP data acquisition and analysis. P.B., L.A.-H. and S.R. wrote the manuscript with input from all authors.

