## Supplementary material for "Principles of mRNA targeting via the Arabidopsis m^6^A-binding protein ECT2": Figure Supplements

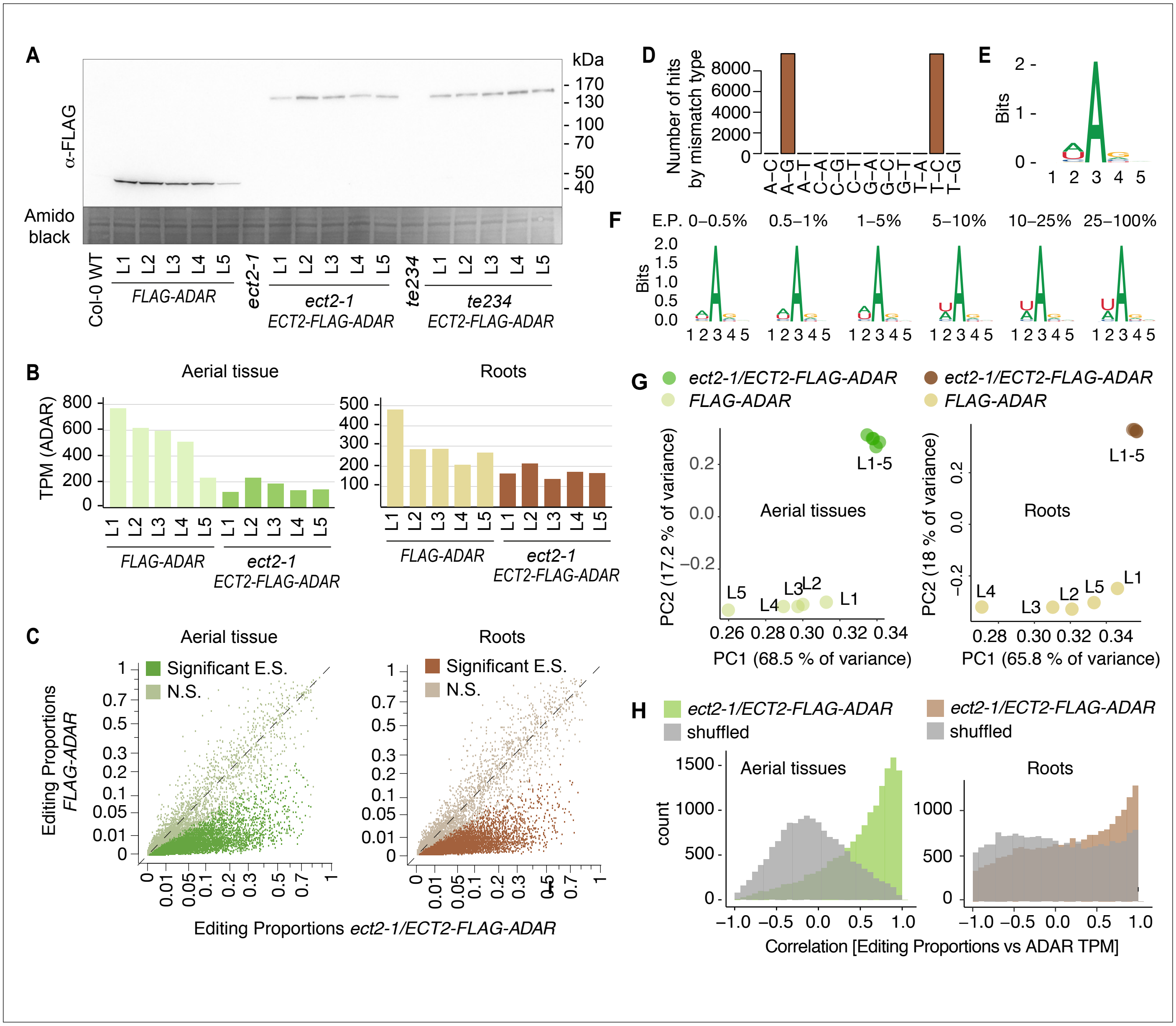


**Figure 1—figure supplement 1. *Drosophila* ADARcd fused to ECT2 can edit target mRNAs *in vivo* in plants (extended data, aerial and root tissues)**. **(A)** Protein expression levels of *ECT2pro:ECT2-FLAG-DmADAR$^{E488Q}$cd-ECT2ter* (*ECT2-FLAG-ADAR*) or *ECT2pro:-FLAG-DmADAR$^{E488Q}$cd-ECT2ter (FLAG-ADAR)* transgenes in 10-day-old seedlings of 5 independent transgenic lines (L1-L5) of each of the genotypes shown in Fig. 1A and their background controls. Amido black staining is used as loading control. **(B)** mRNA expression levels (TPM) of *ECT2-FLAG-ADAR* or *FLAG-ADAR* in dissected apices (aerial tissues) or root tips of the lines used for ECT2-HyperTRIBE (ECT2-HT). **(C)** Scatterplot of the editing proportions (E.P.), defined as G/(A+G), of potential and significant editing sites (E.S.) in aerial and root tissues of *ect2-1/ECT2-FLAG-ADAR* lines compared to the *FLAG-ADAR* controls. Significant sites are highlighted in vivid colours. N.S., not significant. **(D)** Number of significant hits in root samples by mismatch type, after filtering but before removing non-A-G or non-T-C sites. **(E,F)** Consensus motif identified at significant editing sites in roots of *ect2-1/ECT2-FLAG-ADAR* lines, split in groups by editing proportions (E.P.) in F. **(G)** Principal component analysis of editing proportions for significant editing sites. **(H)** Distribution of the correlations between editing proportions and ADAR expression (TPM) for significant editing sites. Background correlations (grey) are based on randomly shuffling ADAR expression for each site. **(A)** Number of hits (significant editing sites) per gene, max or mean editing proportions per gene, and significance of editing sites according to either min or mean -log$_{10}$(-adjusted *p*-value) per gene in ECT2-HT targets split according to their expression levels (in the 5 ECT2-HT control samples).

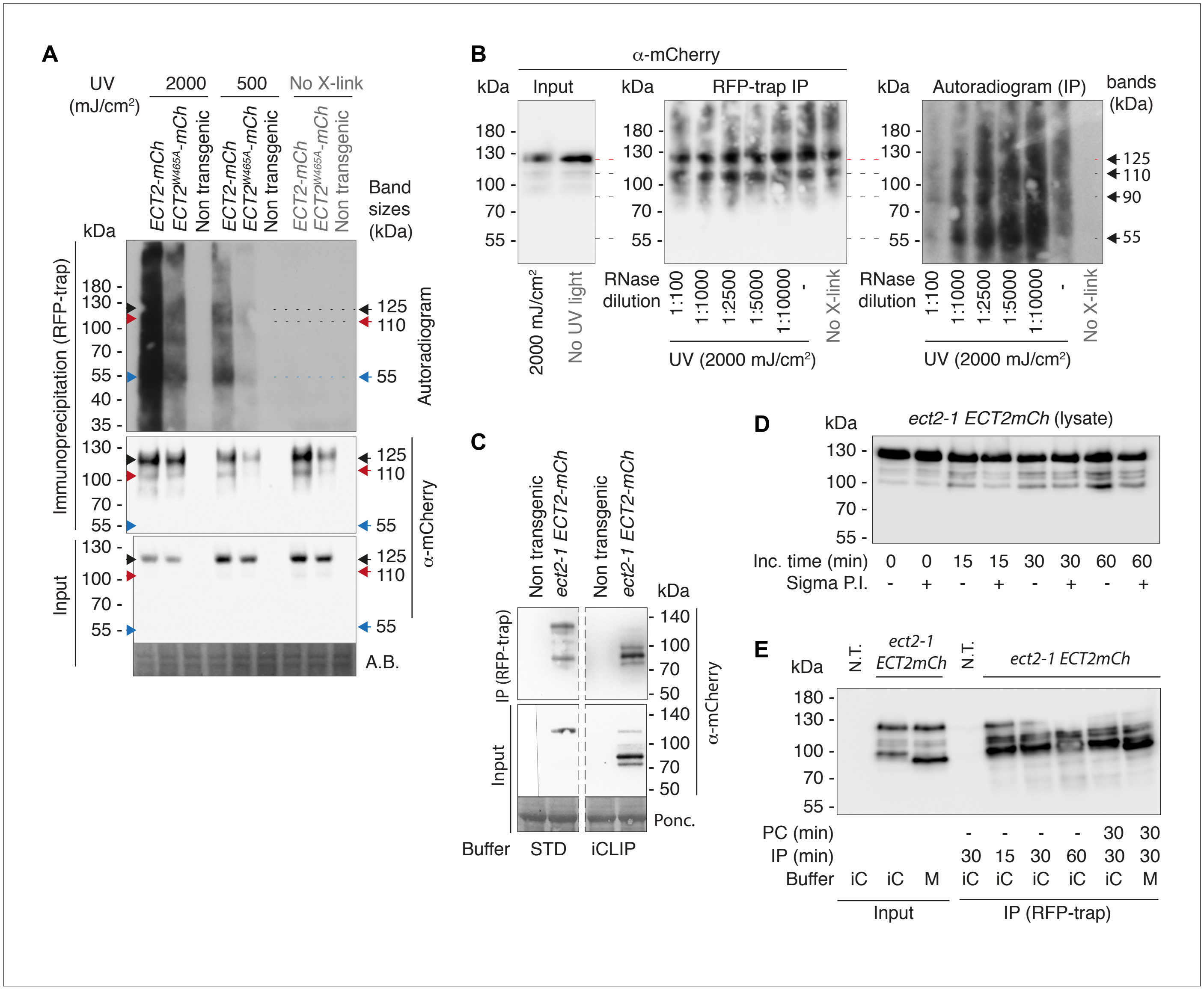


**Figure 3—figure supplement 1. UV-crosslinked RNA co-purifies with ECT2-mCherry in a pattern that depends on the proteolytic cleavage of the ECT2 IDR in the lysate. (A)** Independent repeat of the CLIP experiment in Fig.3B. Sizes corresponding to full length ECT2-mCherry (~125 kDa) (Fig.3C) and the most apparent RNA bands are indicated. **(B)** Same as A using a gradient of increasing concentrations of RNase I. Dilutions refer to the 100 U/µL stock, from which 5 µL were added to 100 µL reactions (e.g. 1:5000 corresponds to the final concentration of 1 U/mL used in all other CLIP experiments and for construction of iCLIP libraries). Note that background in the IP blot is presumably due to lack of cooling during the 3-hour-long SDS-PAGE in this first experiment. **(C)** α-mCherry protein blot of lysates (input) or immuno-purifications (RFP-trap, 1 hour incubation at 4°C either standard IP buffer (STD, [50mM Tris-HCl, pH 7.5, 150mM NaCl, 5mM $MgCl_2$, 10% glycerol, 4 mM DTT, 0.1% Nonidet P-40]) or iCLIP buffer [50 mM Tris-HCl pH 7.5, 150 mM NaCl, 4 mM $MgCl_2$, 5 mM DTT, 1%SDS, 0.25% sodium deoxycholate, 0.25% Igepal], both supplemented only with Roche EDTA-free Protease Inhibitor Cocktail (1 tablet/10 mL). All IP or input lanes have been developed identically on the same membrane. Dashed lines indicate cropping of lanes containing samples irrelevant for this work. Part of the input/non-transgenic/STD-buffer sample lane was accidentally left out when the membrane was developed (the thin line indicates the border of the photograph), but the remaining half lane suggests absence of signal. **(D)** α-mCherry protein blot of cell extracts incubated for increasing amounts of time (at 4°C) in iCLIP buffer supplemented with 4 mM PMSF, 1 tablet/10 mL of Complete Protease Inhibitor Cocktail (Roche), and with or without Sigma Protease Inhibitor Optimized for Plant Extracts (1/30 vol). The progressive, protease inhibitor-sensitive appearance of <125 kDa ECT2-mCherry species is evidence of their generation by proteolysis in the lysate. **(E)** α-mCherry protein blot of lysates (input) or immuno-purifications (RFP-trap) from plants expressing ECT2-mCherry, using different preclearing (PC) and immunoprecipitation (IP) times at 4°C, either in iCLIP buffer (iC) or in a milder (M) variant with ½ amount of detergents, both supplemented with 1 mM PMSF and 1 tablet/10 mL of Complete Protease Inhibitor Cocktail (Roche). Again, the progressive appearance of <125 kDa ECT2-mCherry species with increasing incubation times is seen.

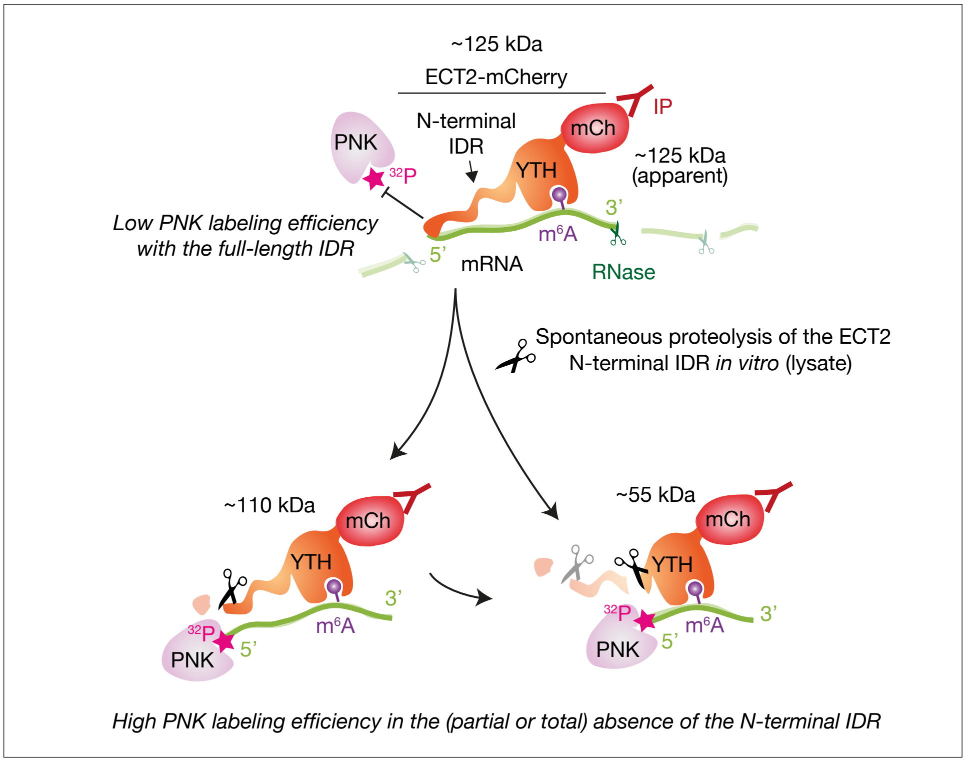


**Figure 3—figure supplement 2. Illustration of RNA-binding properties of ECT2 revealed by CLIP.** Interpretation of the pattern of labelled RNA in CLIP experiments due to the spontaneous proteolysis of the ECT2 IDR in the lysate, and the different labelling efficiency of bound RNA: RNA co-migrating with the most abundant ECT2-mCherry fragment (full-length, ~125 kDa) is barely labeled while the strongest signal appears at ~55 kDa (the size of the YTH domain fused to mCherry), where protein abundance is below the western blot detection limit (Figure 3B,C). This observation suggests limited accessibility of 5’-ends of full length ECT2-bound RNA to polynucleotide kinase (PNK), likely due to binding to the IDR. Supporting this idea, the samples containing full-length protein (’110 kDa band’) required fewer PCR cycles to obtain similar amounts of library DNA and generated more unique reads than their ‘55 kDa band’ conterparts (Figure 3—figure supplements 3,4), indicating that most of the RNA co-purifies with full-length ECT2-mCherry and the stronger intensity of the ‘55 kDa band’ (Figure 3B) is indeed due to differences in labeling efficiency.

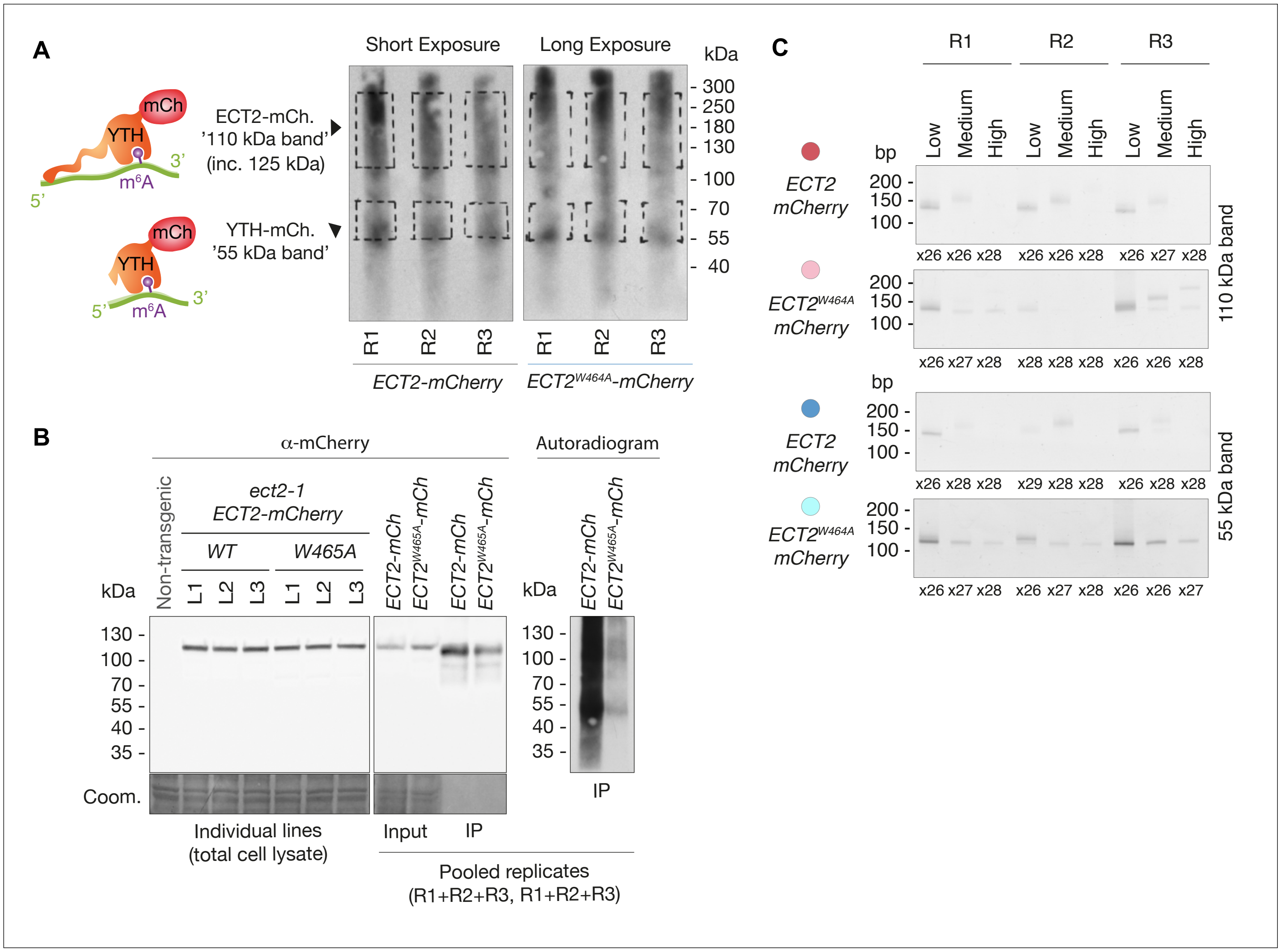


**Figure 3—figure supplement 3. iCLIP Libraries. (A)** Autoradiograms showing labelled RNA co-purified with ECT2-mCherry and $ECT2^{W464A}$-mCherry in the replicates used to generate iCLIP libraries. Handwritten dashed lines on the films were used as guides to excise the corresponding membrane pieces ('110 'Da' and '55 kDa' bands) with a scalpel. The apparently comparable amount of labeled RNA in the two panels, generated from two different membranes and gels run and blotted identically, but separately is a product of different exposure times, which were adjusted in each case for optimal visualization of the RNA. **(B)** Left panel: α-mCherry protein blot from cell extracts of the 3+3 independent lines used for iCLIP. Middle and right panels: α-mCherry protein blot and autoradiogram of the cell extracts (input) and the RFP-trap IPs used to prepare the iCLIP libraries. Because samples for ECT2-mCherry and $ECT2^{W464A}$-mCherry samples were run in separate gels to prevent cross-contamination, pooled aliquots were saved to compare the extent of ECT2 degradation and RNA-binding between them. **(C)** PCR reactions (number of cycles is indicated) used to prepare iCLIP libraries. For each one of the 4 different libraries, the three corresponding PCRs (low, medium and high molecular weight) were combined according to their relative concentrations to compensate for inequalities. Notice that the size of the cDNA insert to be mapped to the genome is expected to be the size of the PCR product minus the length of the P3/P5 Solexa primers and the barcode (128 nt in total) (Huppertz et al., 2014). Therefore, the low molecular weight bands cut at [70–85]-nt on the cDNA gel, with [20–35]-nt-cDNA + 52-nt-primer, generate [145–155]-nt PCR fragments. The low amount of PCR product from the higher molecular weight bands is to be expected, and the not-matching PCR product sizes of the $ECT2^{W464A}$-mCherry control samples are likely a product of the low amount of RNA co-purified with the $m^6A$-binding-deficient ECT2 mutant.

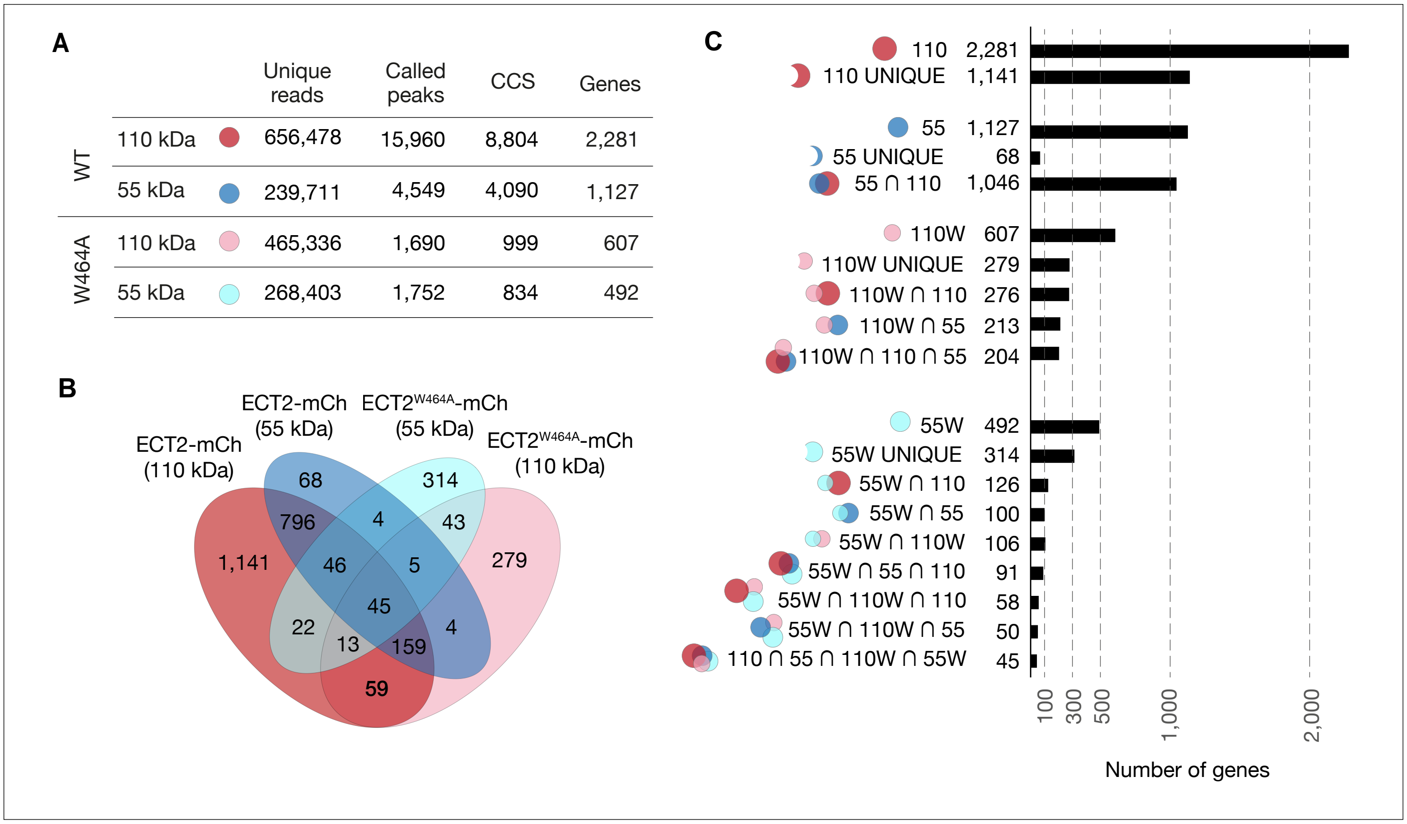


**Figure 3—figure supplement 4**. **Analysis of ECT2 iCLIP Libraries. (A)** Number of valid reads (trimmed reads mapped to the Arabidopsis genome (TAIR10) discarding PCR duplicates), called peaks (= crosslink sites) (Krakau et al., 2017), collapsed crosslink sites (CCS) (peaks with the highest PureCLIP-score within clusters and extended by 4 nt in both directions forming 9 nt wide regions), and target genes, obtained from ECT2-mCherry iCLIP libraries. **(B)** Overlaps between gene sets identified for the 4 ECT2-mCherry iCLIP libraries generated in this study. **(C)** Upset plot displaying gene counts according to different sets defined from the 4 iCLIP libraries.

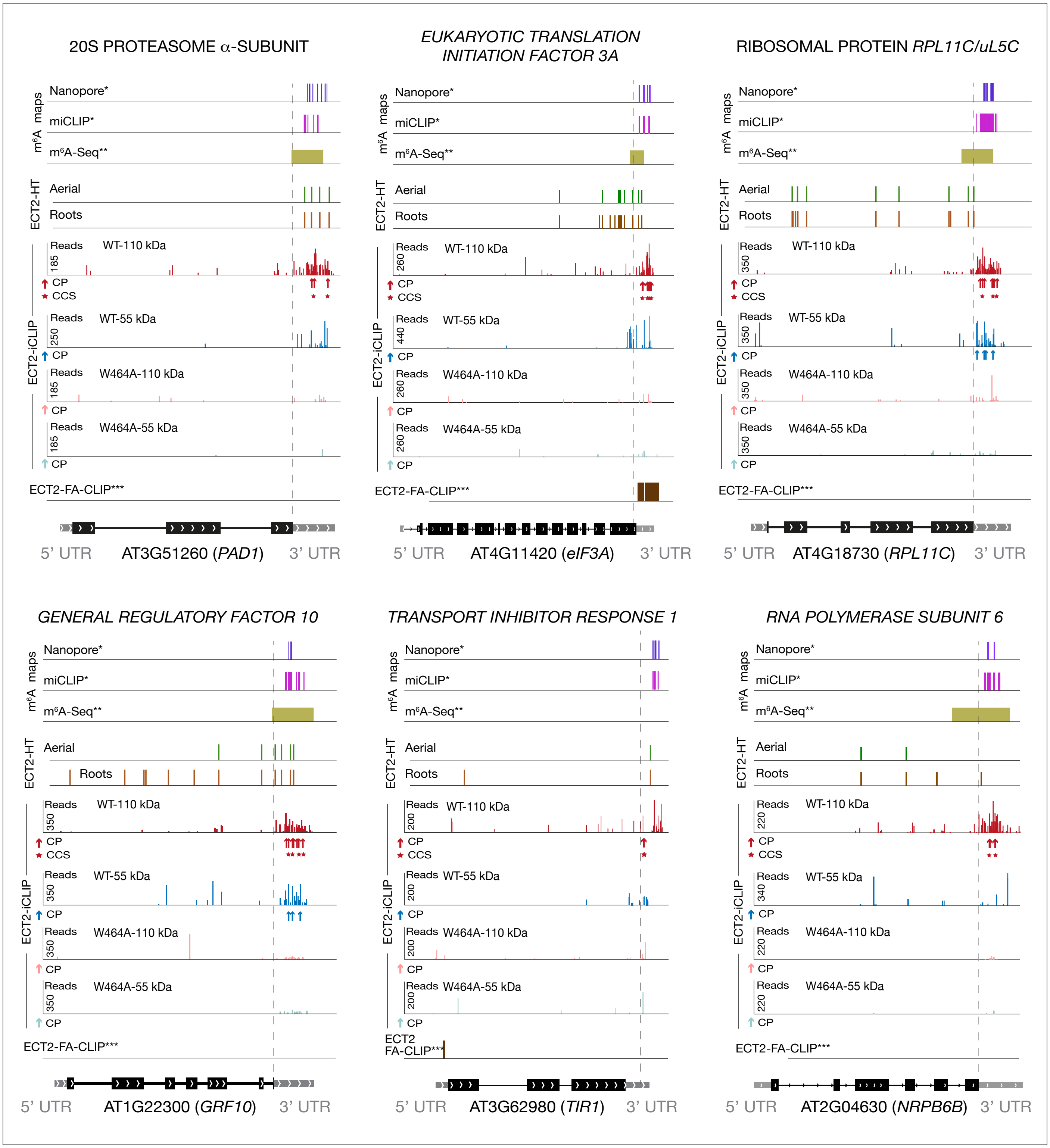


**Figure 4—figure supplement 1. Distribution of m⁶A and ECT2 sites on ECT2 targets**. Representative examples of ECT2 targets showing the distribution of m⁶A sites*, **, ECT2-iCLIP reads and peaks, ECT2-HT edited sites, and FA-CLIP peaks*** along the transcript. CP, called peaks (= crosslink sites); CCS, collapsed crosslink sites. * Parker et al. (2020); ** Shen et al. (2016); *** Wei et al. (2018).

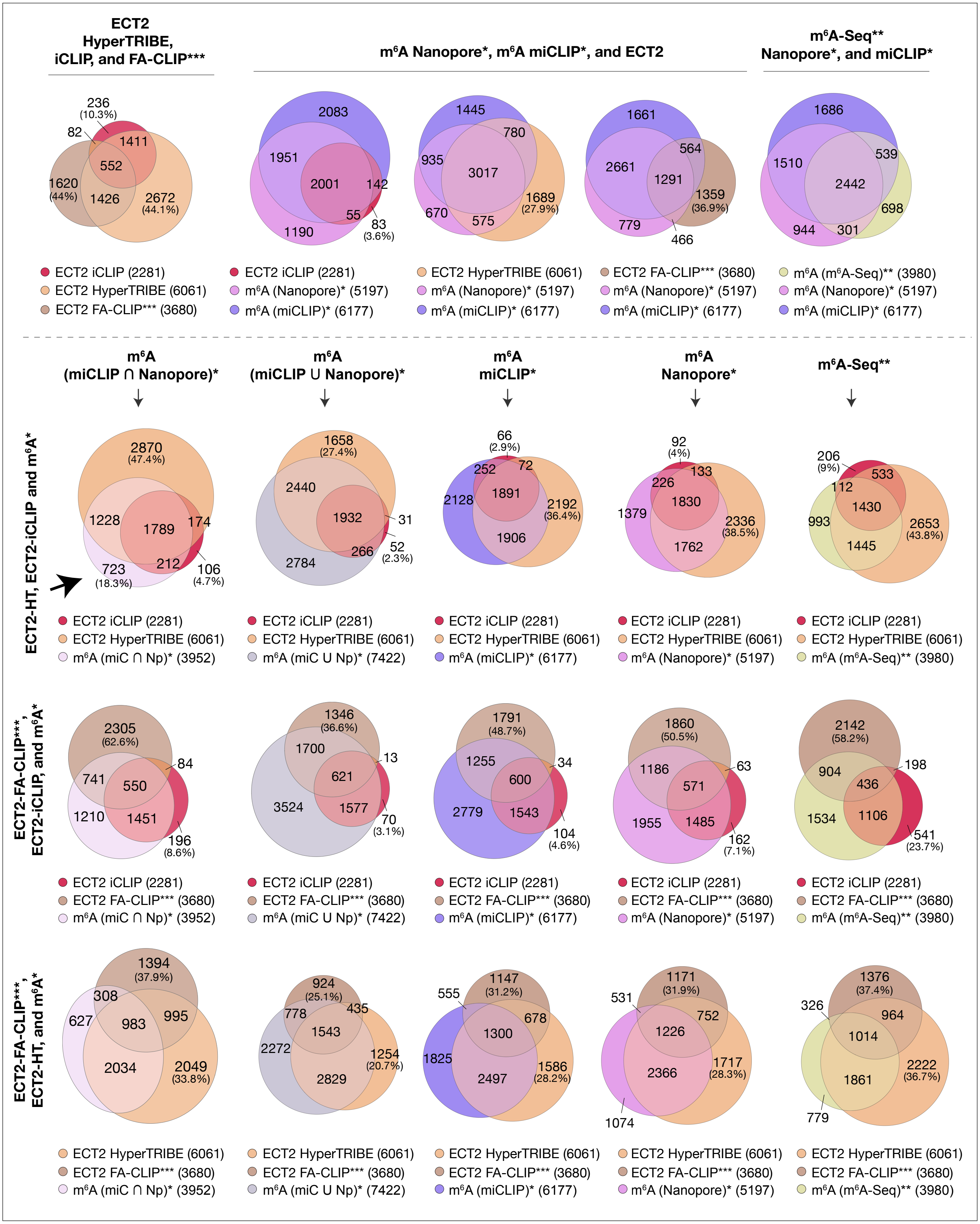


**Figure 4—figure supplement 2. Overlaps between $m^6A$-containing genes and ECT2-targets datasets.** Overlap between genes supported as containing $m^6A$ or ECT2 targets by the different techniques indicated. The ECT2-HT target set includes the sum of targets identified in root and aerial tissues. * Parker et al. (2020); ** Shen et al. (2016); *** Wei et al. (2018).

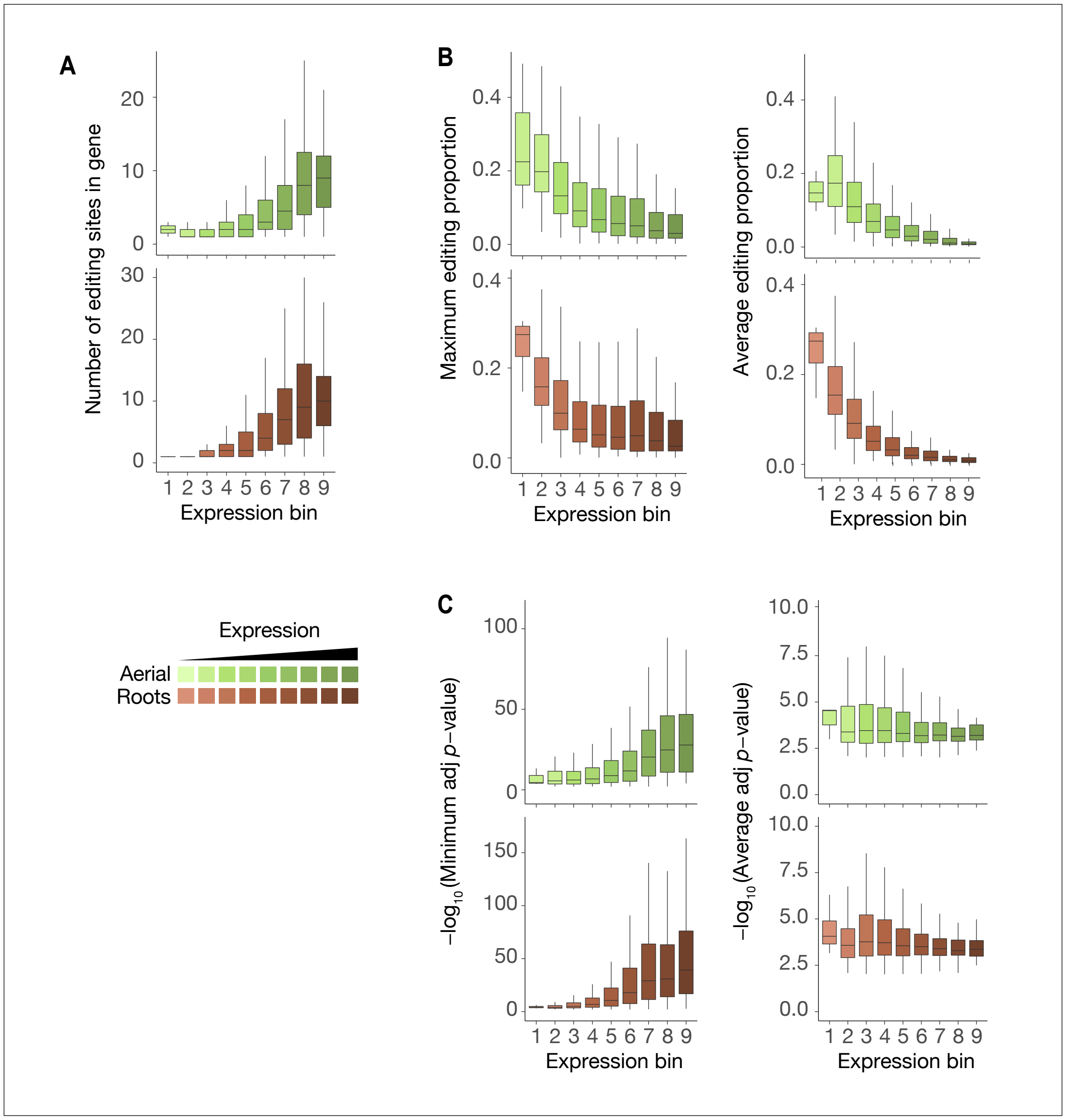


**Figure 4—figure supplement 3. Characteristics of ECT2-HyperTRIBE editing sites relative to target expression levels**. **(A-C)** Number of significant editing sites (A), maximum or average editing proportions (B), and significance of editing sites according to either minimum or average -$\log_{10}$(adjusted *p*-value) per gene (C) in ECT2-HT targets split according to their expression levels (calculated as in Fig. 2B), in both aerial and root rissues. The number of detected editing sites increases with the expression level of the targets (A) while, unexpectedly, the editing proportion decreases (B). This may be caused by dilution, as the ECT2 promoter is active only in highly dividing cells (Arribas-Hernández et al. 2020) while some abundant target mRNAs may be ubiquitously expressed. Nevertheless, the average statistical significance over all edited sites per target mRNA did not change with the expression level (C, right panels), indicating that our HyperTRIBE experimental setup and data analysis identifies targets across a wide range of expression levels including lowly expressed transcripts. This suggests that the significance of differential editing versus a negative control, rather than raw editing proportion or number of editing sites, is the preferred parameter for definition and ranking of targets.

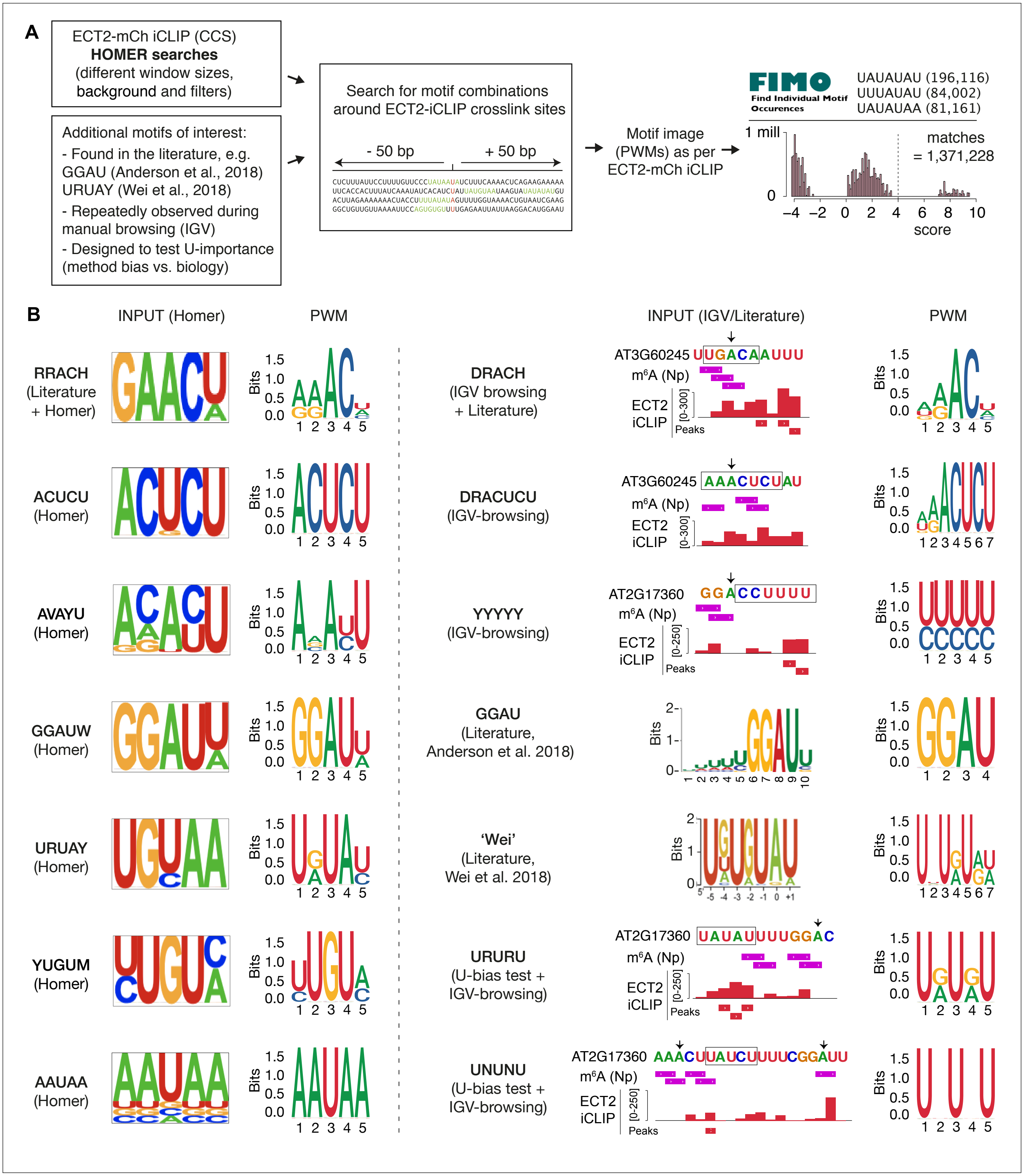


**Figure 5—figure supplement 1. Sources of motifs and generation of position weight matrices. (A)** Relative nucleotide frequencies for motifs of interest with potential sequence redundancies were estimated from windows around ECT2 iCLIP collapsed crosslink sites (CCS) and converted into position weight matrices (PWMs) that were subsequently used in FIMO to scan for motif matches genome wide. Plot illustrates score distribution for the motif derived from Wei et al (2018), and vertical dotted line indicates chosen score cut-off. **(B)** Subset of motifs according to source. Left: examples of motifs inspired by Homer searches (left logo) and subsequent PWM used in the analysis (right logo). Right: examples of further motifs derived from other sources. Logo from the relevant paper is shown if the motif derives from literature, and representative examples of IGV-browser screenshots are shown otherwise (motifs are outlined; arrows mark **A** within DR**A**CH/GG**A**U contexts; Np, Nanopore (Parker et al., 2020). Subsequent PWM is also indicated (right logo).

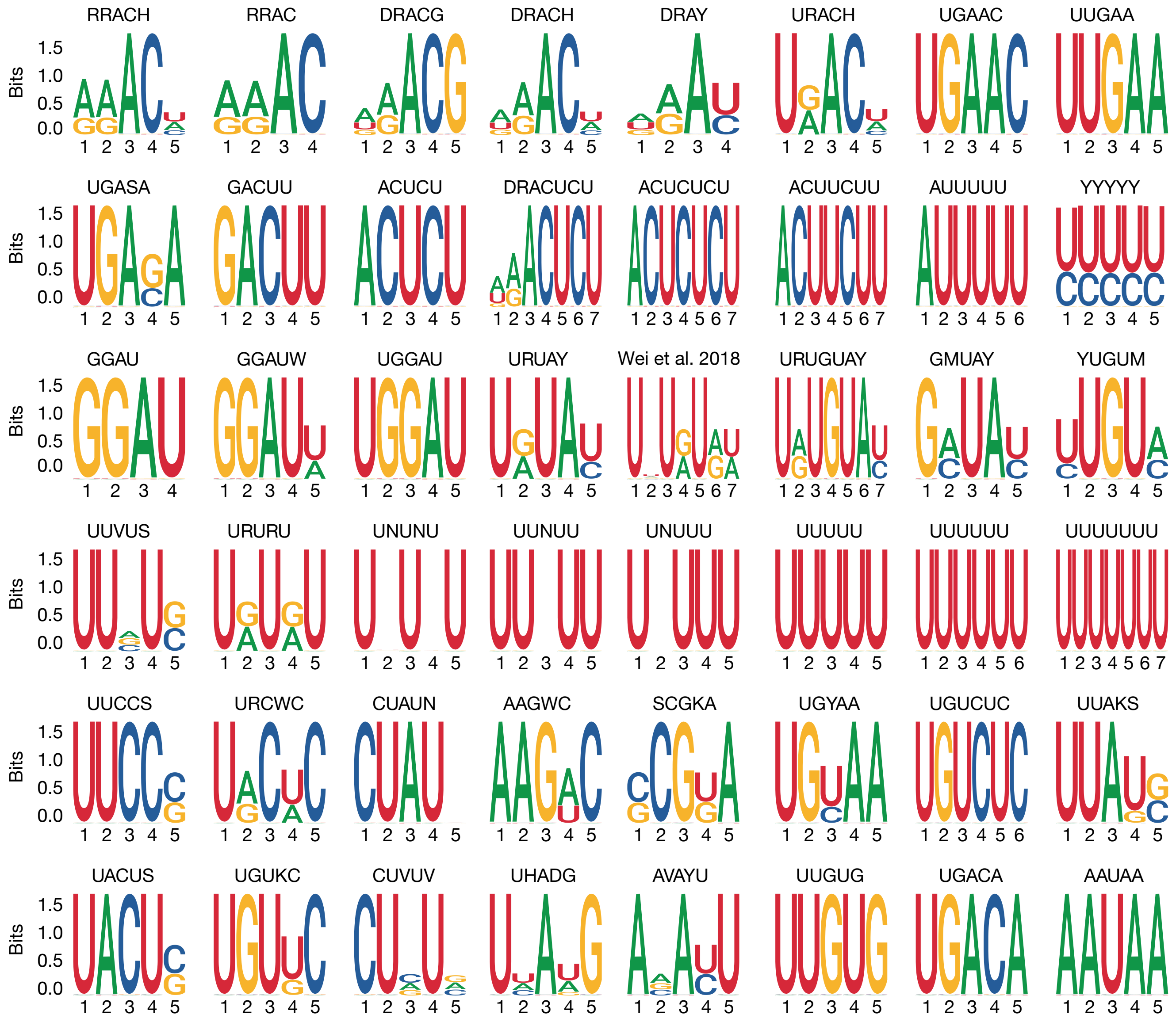


R= A/G  K= G/U  S= C/G  B= C/G/U (not A)  N= A/C/G/U

Y= C/U  M= A/C  W= A/U  D= A/G/U (not C)

H= A/C/U (not G)

V= A/C/G (not U)

**Figure 5—figure supplement 2. Motif logos generated from position weight matrices.** Motif logos represent all of the 48 derived motif position weight matrices considered in the current analysis (see methods and the figure supplement 1 for selection details). UPAC-IUB codes to define multiple nucleotide possibilities in one position are detailed below.

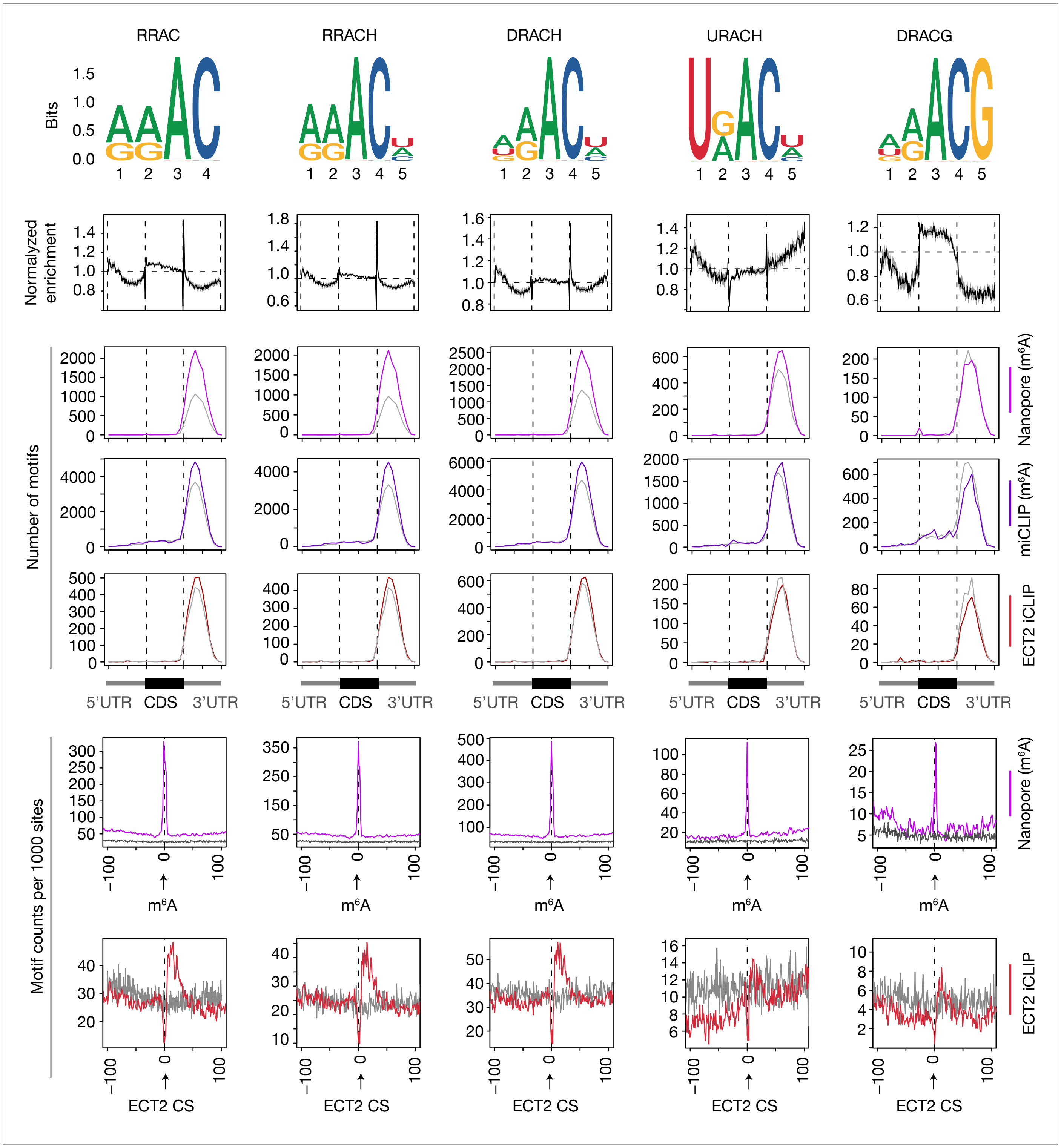


**Figure 5—figure supplement 3. Enrichment of RRACH variants around m⁶A and ECT2 sites.** From top to bottom: (1) motif logos for derived position weight matrices (PWMs); (2) normalized enrichment of motif locations across gene body; (3-5) total number of the relevant motif found at m⁶A-Nanopore* (3), m⁶A-miCLIP* (4) or ECT2-iCLIP (5) sites according to gene body location. Grey lines indicate numbers found in a gene-body location-matched background set of sites of equivalent number; (6-7) distribution of the relevant motif relative to m⁶A-Nanopore* (6) or ECT2–iCLIP (7) sites. Grey lines represent the distribution for the same gene-body location-matched set as derived in the panels above. RRACH shows a slightly higher enrichment over RRAC around Nanopore* sites, that is further increased in the more lenient version DRACH. Accordingly, there is also clear enrichment of URAC. On the contrary, there is no global enrichment of DRACG in Nanopore* datasets along the gene body, and only very modest around Nanopore* sites, highlighting the importance of the final H in DRACH. R=A/G, H=A/C/U, D=A/G/U. * Parker et al. (2020).

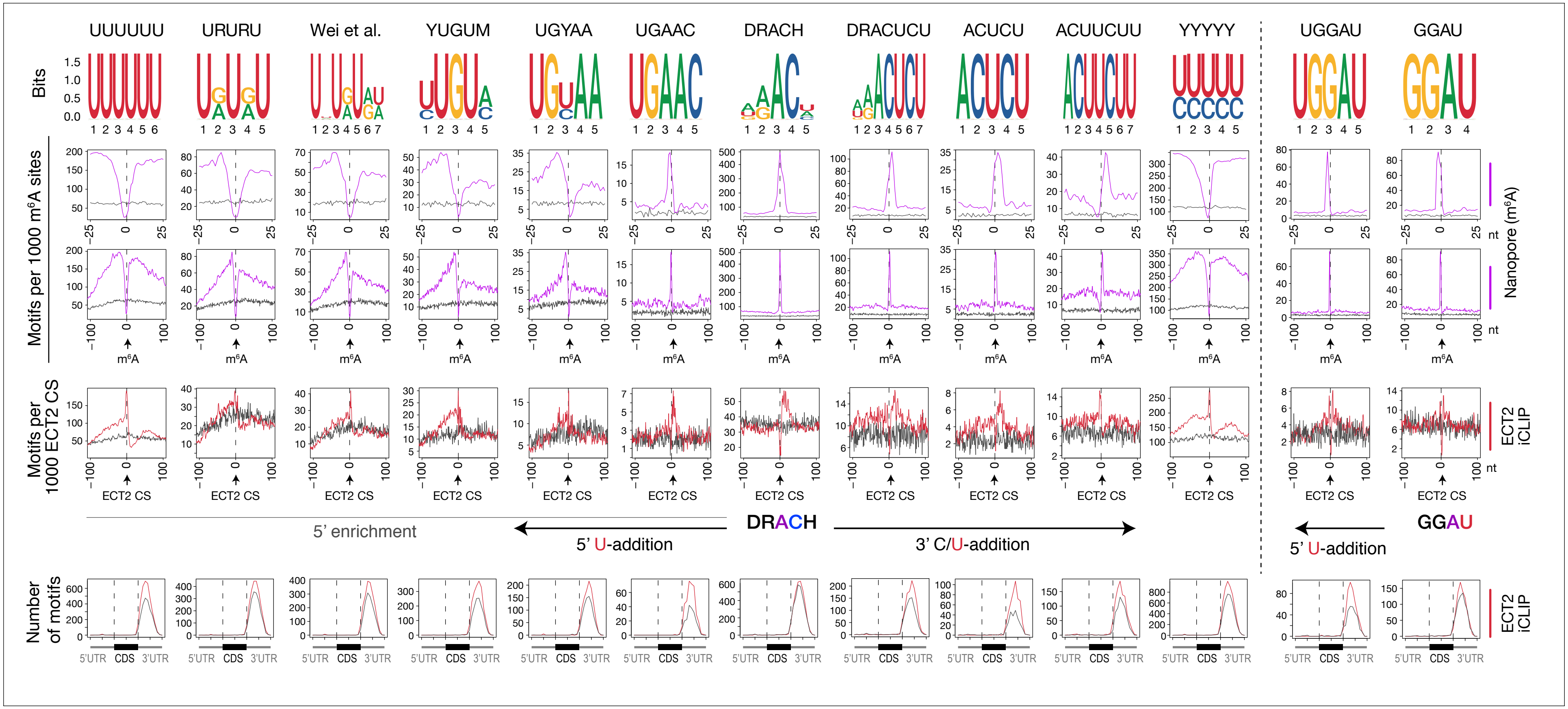


**Figure 5—figure supplement 4. Uridines flanking DRACH result in additional motifs enriched at ECT2 iCLIP sites.** Enrichment of DRACH-like motifs containing additive amounts of U(/Y)s at the flanks until there is only U/Ys. Through the upstream U-additions, transition forms like URURU/UGUAY/YUGUM are included. GGAU with addition of one U upstream is also shown on the right side of the figure. From top panel to bottom: (1) motif logos for derived position weight matrices (PWMs); (2-4) distribution of the relevant motif relative to m⁶A-Nanopore (Parker et al., 2020) (2,3) or ECT2–iCLIP crosslink sites (CS) (4). Grey lines represent the distribution for gene-body location-matched background set of sites of equivalent number; (5) total number of the relevant motif found at ECT2-iCLIP sites according to gene body location. Grey lines indicate numbers found in the same gene-body location-matched background set.

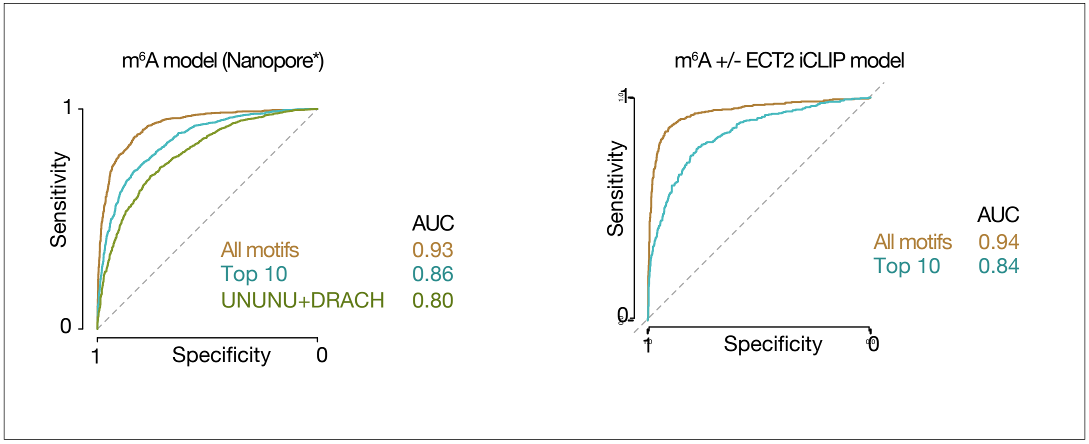


**Figure 6—figure supplement 1. Model performance ROC curves for distinguishing sequence preferences of either m⁶A or ECT2-bound sites. (A,B)** Receiver operating characteristic (ROC) curves showing model performance of random forest (gradient boosting) trained models, for distinguishing either m⁶A Nanopore* sites from their respective random location-matched background sites (A), or m⁶A sites with and without ECT2 iCLIP crosslink sites (B), with overall performance represented by area under the curve (AUC) values. 'All motifs' refers to the full set of 48 motifs (at, upstream or downstream), 'Top 10' to the top 10 most important features from the full model, and "UNUNU+DRACH" includes features derived only from UNUNU or DRACH motifs. *Parker et al. (2020).

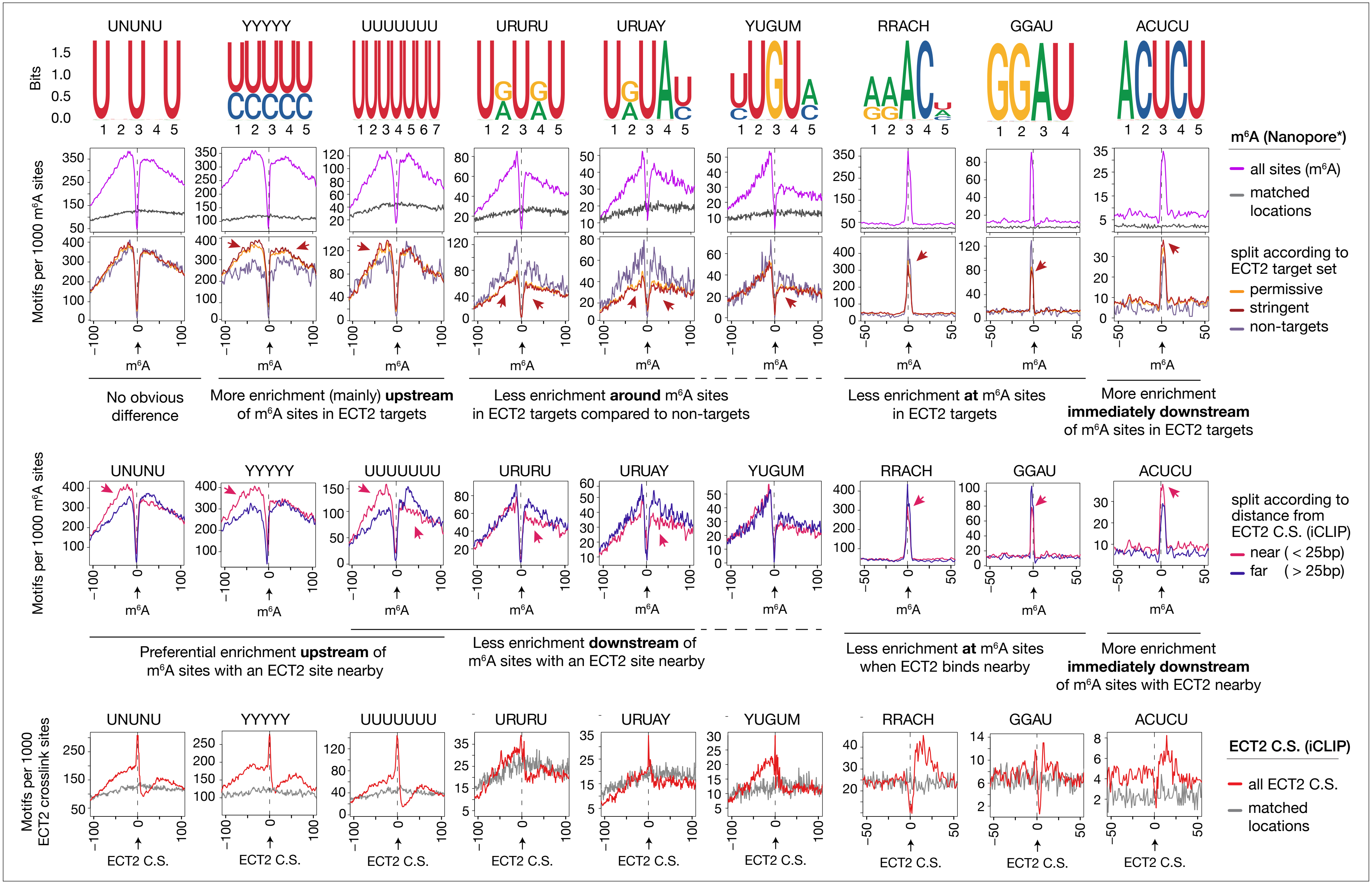


**Figure 7—figure supplement 1. Motif preferences around m6A sites according to ECT2 binding.** From top to bottom: (1) motif logos for derived position weight matrices (PWMs); (2) distance-based enrichment of motifs at and around m6A-Nanopore* sites, plotted as motif counts per 1000 m6A sites (purple lines). Grey lines indicate the enrichment in a location-matched background set as in Figure 5D; (3) same as in 2 with sites split according to whether they sit on ECT2 targets; (4) same as in 2 with sites split according to the distance from the nearest ECT2 crosslink site (for ECT2-iCLIP targets only); (5) motif counts per 1000 iCLIP binding sites, as a function of distance from the iCLIP position, showing all sites against matched background sites (grey lines). Motifs are ordered according to the different relative enrichment upstream or downstream m6A sites according to ECT2 binding (see text below panels 3 and 4). * Parker et al. (2020).

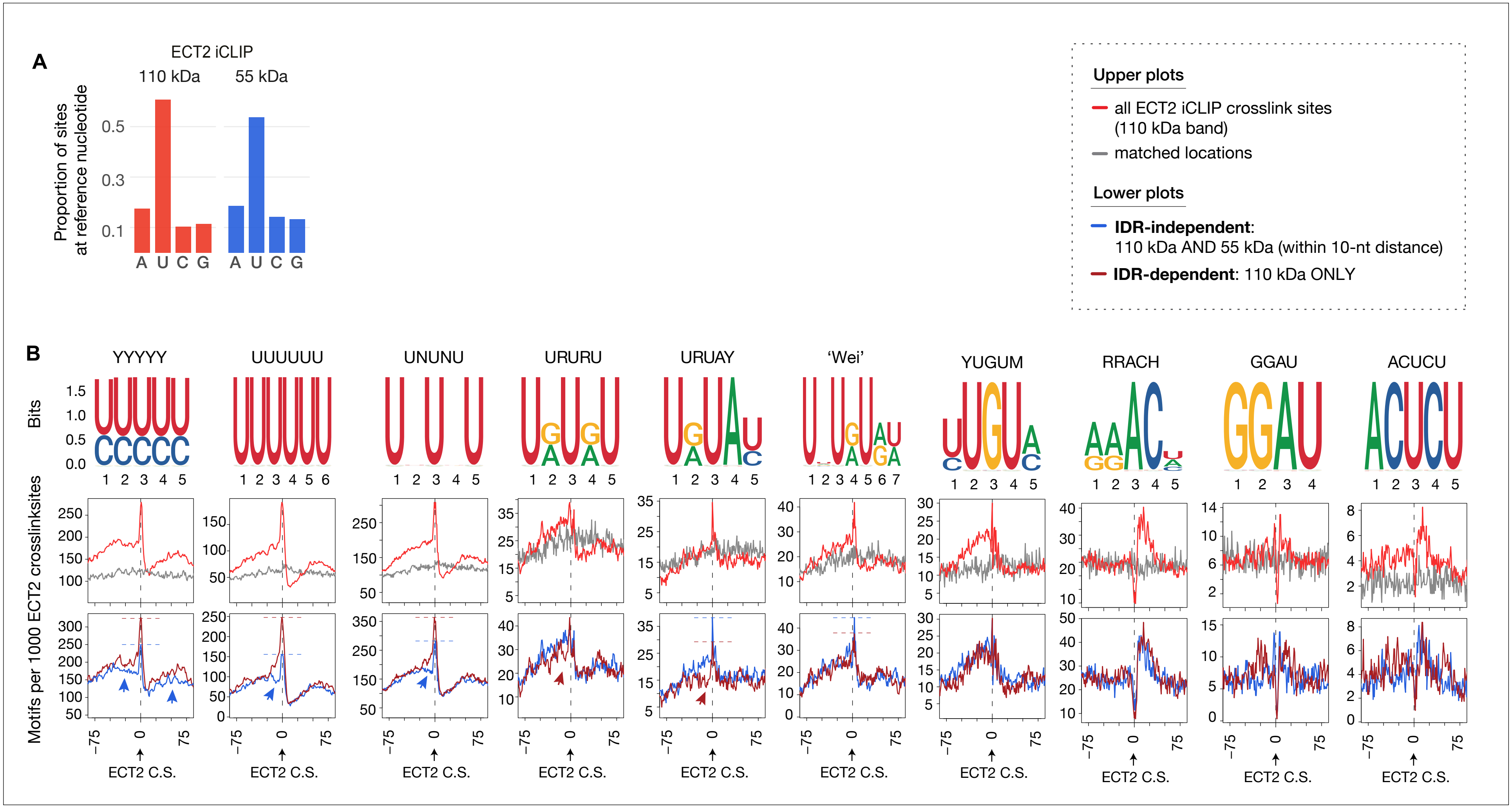


**Figure 7—figure supplement 2. Dependency of the ECT2 IDR for motif enrichment. (A)** Proportion of ECT2 iCLIP sites at each nucleotide for the 110 kDa and 55 kDa bands. **(B)** From top to bottom: (1) motif logos for derived position weight matrices (PWMs); (2) distribution of motifs per 1000 ECT2 iCLIP crosslink sites (red) or matched-locations background (grey), +/- 75 nt of the site; (3) Motifs per 1000 ECT2 iCLIP crosslink sites, split according to whether they are found in libraries from both 110 kDa and 55 kDa bands ('IDR-independent'), or exclusively (distance > 10 nt) in the 110 kDa band ('IDR-dependent').
