## Supplemental Methods for "Principles of mRNA targeting via the Arabidopsis m^6^A-binding protein ECT2"

*Plant growth conditions*

Seeds were surface-sterilized by 2-min incubation in 70% EtOH plus 10 min in [1.5% NaOCl, 0.05% Tween-20], two $H_2O$ washes, and 2-5 days of stratification at 4°C in darkness. Plates with T2 seedlings for HyperTRIBE were supplemented with glufosinate ammonium (Fluka) (7.5 mg/L) to select plants expressing the ADAR-containing transgenes. To assess phenotypes of adult plants, ~8-day-old seedlings were transferred from horizontal MS plates (4.4 g/L MS, 10 g/L sucrose, 8 g/L agar; pH 5.7) to soil and maintained in Percival incubators under 16 hr light/8 hr dark cycles, 21°C-day/18°C-night temperature, and ~100 µmol $m^{-2}$ $s^{-1}$ light intensity. We used Philips fluorescent tubes TL-D 90 De Luxe 36W as light source.

*Cloning*

To build the constructs for HyperTRIBE by USER cloning (Bitinaite and Nichols 2009), fragments containing *ECT2* gDNA sequences were amplified by PCR (KAPA HiFi Hotstart Uracil+ ReadyMix, Roche) from plasmids previously generated in our lab (Arribas-Hernández et al. 2018). The *FLAG-DmADAR$^{E488Q}$cd* fragment was produced in the same way using a pGEM-T Easy (Promega) plasmid containing FLAG-*DmADAR$^{E488Q}$cd* as template, previously subcloned to introduce the E488Q hyperactive mutation by site-directed mutagenesis (QuickChange, Agilent Technologies) with primers LA729-LA730 (Phusion HF DNA Polymerase, NEB). The E488Q mutation was detected by *NlaIII* (ThermoFisher) digestion of the PCR reaction (DreamTaq, ThermoFisher) obtained with primers LA660-LA735. Of note, the *FLAG* and *DmADARcd* sequences had been previously glued together by USER-cloning to produce *AGO1pro:FLAG-DmADARcd-AGO1ter* in pCAMBIA3300U for unrelated purposes (unpublished work), and subsequently amplified together by PCR with primers LA696-615 for introduction into pGEM-T-Easy. To build *AGO1pro:FLAG-DmADARcd-AGO1ter* in the first place, the catalytic domain of the ADAR deaminase isoform N (Y268-E669) was amplified from cDNA of *Drosophila melanogaster* Canton-S wild type flies and larvae with USER primers MVUSER12-22. The rest of the fragments were amplified from *pCAMBIA3300U AGO1pro:FLAG-AGO1-AGO1ter* (Arribas-Hernández et al. 2016) with primers MVUSER1-11 and MVUSER23-6.

USER primers to amplify all fragments were designed to create overhangs compatible with either the *PacI* USER cassette present in the pCAMBIA3300U plasmid (see Methods in the main text), or the flanking sequences of the neighboring fragments.

All primer sequences, their combinations to produce PCR fragments and the arrangement of the fragments for USER cloning can be found in Table 1 of the Supplemental Material.

Kanamycin-resistant colonies were analyzed by restriction digestion and sequencing prior introduction of the plasmids in Agrobacteria (strain GV3101) for plant transformation.

*Western blotting*

Protein extraction from 10-day-old seedlings and western blotting with FLAG, HA and mCherry antibodies were done as previously described (Arribas-Hernández et al. 2018). Loading was documented by amido black (A.B.), Coomassie or Ponceau staining of the total protein on the membrane.

*RNA extraction and library preparation for HyperTRIBE*

The tissue was flash-frozen in liquid nitrogen and ground into a fine powder using liquid nitrogen-cooled adaptors in a tissue homogenizer. For RNA extraction, we added 1 mL of TRI Reagent (Sigma) to the frozen tissue (<100 mg), mixed quickly by vortexing, added 0.2 mL of chloroform, and separated the 2 resulting phases by vigorous shaking and 10 min centrifugation at 4ºC. The RNA was then precipitated from the aqueous phase for 30 min at room temperature with 1 volume of isopropanol. RNA pellets were solubilized in 300 µL of $H_2O$ to remove polysaccharides through a mild precipitation by addition of 1/10 vol. 99% EtOH and 1/30 vol. of [3M NaOAc pH 5.2] and incubation on ice for 30 min. After 15 min of full-speed centrifugation at 4ºC to pellet polysaccharides, we re-precipitated the RNA from the supernatant with 2,5x vol. 99% EtOH and 1/10 vol. of [3M NaOAc pH 5.2], washed the pellet 2 times with 70% EtOH, and resuspend in 20-40 µL of $H_2O$. This highly pure total RNA was then used to produce mRNA libraries through enrichment of mRNA with oligo(dT) beads (18-mers), random fragmentation, cDNA synthesis with random hexamers, custom second-strand synthesis (Illumina), terminal repair, A-ligation and sequencing adaptor ligation, size selection (250~300 bp insert) and PCR enrichment. The libraries were prepared and sequenced (Illumina PE150, Q30≥80%) as a service from Novogene.

*CLIP experiments and iCLIP library preparation*

Crosslinked plant tissues (see details below) were finely ground in liquid nitrogen with mortar and pestle, homogenized in iCLIP buffer [50 mM Tris-HCl pH 7.5, 150 mM NaCl, 4 mM $MgCl_2$, 5 mM DTT, 1%SDS, 0.25% sodium deoxycholate, 0.25% Igepal] supplemented with protease inhibitors (4 mM PMSF, 1 tablet/10 mL of Complete Protease Inhibitor Cocktail (Roche), and 1/30 vol. of Protease Inhibitor Optimized for Plant Extracts (Sigma P9599)), and cleared by centrifugation and filtration (0.45 µm pore) of the supernatant. RNP-complexes were then immunopurified with beads coupled to anti-RFP nanobodies (ChromoTek RFP-Trap in our case) for 1 hour at 4ºC under constant rotation. In particular, we used 20 µL of beads for 4 g of tissue in 6 mL of iCLIP buffer for every replicate. After thorough washes with RIP-Wash Buffer [2M urea, 50 mM Tris-HCl pH 7.5, 500 mM NaCl, 4 mM $MgCl_2$, 2 mM DTT, 1% SDS, 0.5% sodium deoxycholate, 0.5% Igepal], RNP-complexes attached to the beads were subjected to treatment with DNase (Turbo DNase (Ambion), 4U/100 µL) and RNase I (Ambion, 1 U/mL) at 37°C for 10 min, dephosphorylation of RNA 3' ends (PNK (ThermoFisher) in pH 6.5 buffer), and 3' RNA linker ligation (L3-App linker (Huppertz et al. 2014) and NEB HC RNA Ligase) at 16°C overnight. RNA was radioactively labeled at the 5' end by PNK-mediated phosphorylation using γ-$^{32}$P-ATP (20 min at 37°C). The labeled RNP complexes were subjected to SDS-PAGE and blotting on a nitrocellulose membrane (Protran BA-85). Pieces of membrane containing a size-range of RNA species bound to the protein (a smear above the expected molecular weight localized by autoradiography) were excised and subjected to proteolysis (200 µg of Proteinase K (Roche) in 200 uL of PK buffer [100 mM Tris-HCl pH 7.4, 50 mM NaCl, 10 mM EDTA] for 20 min at 37°C) to release RNA bound to small peptides. The RNA was then purified with TRI-Reagent (Sigma) and used to prepare sequencing libraries through the following steps: Reverse transcription (Superscript III, Invitrogen) using a two-part cleavable DNA adapter complementary to the 3' RNA linker as primer, gel purification and size selection of cDNA (high, 120-200 nt; medium, 85-120 nt; low, 70-85 nt), circularization (CircLigase II (Epicentre), relinearization (BamHI), and PCR amplification. All steps were performed as described by Huppertz et al. (2014), and the amount of cycles in the final PCR was optimized to the amount of cDNA in each sample.

Notice that we introduced a few modifications in the original protocol (Köster and Staiger 2020) to account for: 1) Low abundance of ECT2 compared to AtGRP7. To obtain enough RNA, we increased the crosslinking energy and irradiated 12-day-old seedlings with 2000 mJ/cm$^2$ of 254 nm UV light, harvesting roots and shoots (4 g of tissue per replicate) to maximize the amount of purified ECT2-mCherry. 2) ECT2 sensitivity to proteolysis. We did not pre-clear the lysates to reduce the incubation time, and we used high amounts of protease inhibitors during immunoprecipitation. 3) High molecular weight of ECT2-mCherry. Due to the size of the protein, we required longer electrophoresis time and cooling (3 hours at 180 V with the tank on ice). 4) Different RNA-binding capacity of ECT2. Based on trials, we decided to adjust the RNase I treatment to 1 U/mL, incubating for 10 min at 37°C (5 µL of RNase I (Ambion, 100 U/µL pre-diluted 1:5000) in 100µL).

Of note, the conditions indicated here were specifically used for library preparation. Although we used the same conditions as default for CLIP experiments to assess ECT2 RNA-binding capacity, ECT2 sensitivity to proteolysis and ECT2-bound RNA sensitivity to RNase treatment, variations in buffer composition, incubation time, concentration of protease inhibitors and/or RNase I are specified in the corresponding figure legends where necessary.

*HyperTRIBE data analysis*

Significant differentially edited sites between *ECT2-FLAG-ADAR* (fusion) and *FLAG-ADAR* (control) samples for ECT2 HyperTRIBE (ECT2-HT) were called according to our hyperTRIBE**R** pipeline (https://github.com/sarah-ku/hyperTRIBER). First, reads were trimmed using *trimmomatic* (Bolger et al. 2014) and mapped to the Arabidopsis genome (TAIR10) using STAR (Dobin et al. 2013), according to parameters suggested in a previous HyperTRIBE analysis (Xu et al. 2018). A custom perl script based on SAMtools *mpileup* (Li et al. 2009) returned base counts for all positions where there is a mismatch from the reference in at least one sample. For running the hyperTRIBE**R** analysis pipeline, we specified that any tested position must have a putative edit in at least 4 of the 5 replicates in the fusion samples (3 of 4 in the case of roots since one of the *ECT2-FLAG-ADAR* samples, "L3", was deemed as low quality and subsequently removed from the significance calling pipeline). Significant hits (adjusted $p$-value < 0.01 and $\log_2$FC > 1) were further filtered as follows: 1) hits which did not correspond to an A-to-G change (or a T-to-C change for the negative strand) 2) hits which were likely SNPs arising specifically in either the *ECT2-FLAG-ADAR* or *FLAG-ADAR* line manifesting in an editing proportion at or close to 1, and 3) hits where the coverage of tags at the edit base over the *ECT2-FLAG-ADAR* were fewer than 10 reads.

All HyperTRIBE samples were quantified using Salmon (Patro et al. 2017), with appropriate settings for pair-end sequencing and non-stranded library set-up and based on the transcriptome for Araport11 (Cheng et al. 2017) with manual addition of the *FLAG-ADAR* sequence.

Annotations pipeline from hyperTRIBE**R**: editing proportions were calculated as G/(A+G) (alternatively C/(U+C) for the negative strand) for all significant sites, averaged over all samples, separately for the *ECT2-FLAG-ADAR* and *FLAG-ADAR* samples. Significant sites were annotated to genes from Araport11, prioritizing the gene with the highest expression in the given tissue in the case of multiple possibilities. Possible transcripts were subsequently ordered by expression along with gene-body location the position annotates to along the transcript (5'UTR, CDS, 3'UTR). Expression TPM for HyperTRIBE samples was quantified using Salmon (Patro et al. 2017), generated based on the Araport11 transcriptome and annotated TPM in the pipeline was based on averages from *FLAG-ADAR* samples only, respectively for aerial tissues or roots HyperTRIBE results.

Principal component analysis was carried out on the raw editing proportions per sample, for all sites with significant evidence of editing.

Comparison of sites between aerial tissues and roots: genes defined as commonly expressed in both aerial tissues and roots were considered in all gene-based comparisons. For significant editing site-based comparison, we directly compared sites that were common and significant to both.

Correlations with *FLAG-ADAR*: Transcripts per million (TPM) mapping to *FLAG-ADAR* were extracted from quantifications from Salmon (Patro et al. 2017) and correlated with the raw editing proportions per sample, separately for the fusion and control samples. Background correlation estimates were calculated by first scrambling the order of the *FLAG-ADAR* TPM vector.

Identified motif for significant ECT2-HT sites. All sequences were derived from TAIR10 using the R package *rtracklayer* (Lawrence et al. 2009), for bases at and -/+ 2 nt of the significant editing positions, in either aerial tissues or roots. A matrix of nt frequencies (A, C, G or U) was generated and the R package *ggseqlogo* (Wagih 2017) was used to generate the final motif.

Expression binning: $\log_2$(TPM+1) values for all expressed genes in either aerial tissues, roots or combined were split into 9 bins of increasing expression, using the cut() function from the R package *Hmisc* (https://github.com/harrelfe/Hmisc/). For the proportion of genes in every expression bin, annotated genes in each set ($m^6A$ sets (Parker et al. 2020), ECT2 HT/iCLIP-targets or non-targets, and ECT2 FA-CLIP (Wei et al. 2018)) we calculated the proportion of genes falling into each expression bin, out of the total number of genes in that bin. To demonstrate expression biases in unsupported ECT2-HT target genes, the genes were further split according to whether or not they had support from $m^6A$ (Nanopore, miCLIP (Parker et al. 2020) and $m^6A$-seq (Shen et al. 2016)).

Comparison with single cell data – the expression matrix based on a total of 4727 individual cells from scRNA-seq in roots was downloaded from Denyer et al. (2019). To estimate the relationship between co-expression of target genes with ECT2 and their average editing proportions, the expression matrix was used to calculate co-expression for each target gene as follows:

(# cells expressing ECT2 AND target gene) / (# cells expressing target gene)

These proportions were then split into groups and plotted against the maximum editing proportions from HyperTRIBE in the containing genes.

*Analysis of publicly available data*

Single nucleotide resolution locations of $m^6A$ sites defined by either Nanopore sequencing (by comparing samples containing a *vir-1* mutation and wild type samples) or miCLIP were download from Parker et al. (2020). Intervals defining $m^6A$ site locations based on $m^6A$-seq were downloaded from Shen et al. (2016), and intervals defining locations of ECT2 bound sites as determined by FA-CLIP were download from Wei et al. (2018). All sets were annotated using the HyperTRIBE annotation pipeline, based on genes and transcripts from Araport11, for consistency with HyperTRIBE and ECT2-iCLIP gene annotations, specifying the center of the intervals for $m^6A$-seq and FA-CLIP and using the averaged TPM for transcripts from *FLAG-ADAR* samples from ECT2-

HT for relative transcript quantifications.

*iCLIP data analysis and peak calling*

Sequenced reads from all samples were investigated after each processing step with *fastqc* 0.11.5 (https://www.bioinformatics.babraham.ac.uk/projects/fastqc/). Adapters at the 3' end were trimmed using *cutadapt* version 1.16 (Martin 2011). The demultiplexing of the samples was performed using *flexbar* 3.4.0 with the -bk parameter to conserve the barcode information for further steps (Roehr et al. 2017). Reads with a length below 24 nucleotides were discarded. Barcodes were trimmed and saved to the *read_id* field. Processed reads were mapped to the TAIR10 genome with STAR version 2.6.0a allowing a maximum of 2 mismatches and soft clipping only at 3' end (Dobin et al. 2013). PCR duplicates were removed by grouping the reads by their mapping start position. Reads with the identical start position and random barcode were removed from the samples (Python3 and pybedtools). The peak calling of uniquely mapped reads was done using PureCLIP 1.0.4, choosing the second peak-shape option to allow more broader peaks to be called (Krakau et al. 2017).

For consistency with the ECT2-HT datasets, the ECT2-iCLIP datasets were annotated using to the hyperTRIBE**R** annotation, using quantifications based on the average of roots and aerial tissues from *FLAG-ADAR* samples in ECT2-HT (to reflect that the ECT2-iCLIP data is based on whole seedlings).

To calculate the proportion of sites falling at each nucleotide, nucleotide sequences from the reference genome were first obtained from site coordinates for ECT2 iCLIP/m$^6$A-Nanopore/ m$^6$A-miCLIP using R packages *GenomicRanges* (Lawrence et al. 2013) and *rtracklayer* (Lawrence et al., 2009) and nucleotide proportions were plotted using ggplot2 (https://ggplot2.tidyverse.org).

*Motif discovery*

To remove redundancy after the peak calling, directly adjacent peaks (crosslink sites) were grouped together and only the peak with the highest pureCLIP score (dominant) was kept. The called peak position (1 nt resolution) was extended by 4nt up- and downstream to define a 'collapsed crosslink site' (CSS) with length 9nt. The center position marks the dominant called peak. The extension of the peak positions was computed using *bedtools* version 2.27.1 (Quinlan and Hall 2010; Dale et al. 2011).

*Motif enrichment and co-occurrence*

Compiling final list of motifs: motifs found to be significant in Homer (Heinz et al. 2010) searches, based on varied window sizes, settings and backgrounds, as well as centering on either iCLIP binding site-assigned peaks and peaks from Nanopore (Parker et al. 2020), were collated manually. A range of motifs closely related to the consensus motif RRACH (RACH, DRAY, DRACH, URACH, DRACG) were also added to the list, as well as various combinations of U-rich sequence (UUUUU, UNUNU, etc.). Furthermore, motifs found to be of interest in scientific literature were also included, for example URUGUAY/URUAY (Wei et al. 2018), GGAU/GGAUW (Anderson et al. 2018), as well as extra motifs which appeared of potential interest from manually browsing the sequence in the vicinity of iCLIP peaks (YYYYY, DRACUCU, and others, see Supplemental Fig. 7). This resulted in a final list of 48 motifs for further analysis (Supplemental Fig. 8).

Generating position weight matrices and FIMO (Grant et al. 2011): Regions of sequence around ECT2 iCLIP peaks were scanned for a given, potentially redundant, motif patterns. For each motif, the relative frequencies of nucleotides at each position were used to construct a custom position weight matrix, using the formula PWMb,j = $\log_2$ [p(b,j)/p(b)], where p(b) is the background frequency of each nucleotide (see further down), p(b,i) is the frequency of the nucleotides in each position j. We also included an extra small frequency count in the calculation to account for potential uncertainty in redundancies. Logos for all motifs were generated using the Rpackages ggplot2 (https://ggplot2.tidyverse.org) and *ggseqlogo* (refs). The calculated position weight matrices were run through FIMO 5.1.1 (Grant et al. 2011), specifying background letter frequencies (A: 0.273, C: 0.165, G: 0.173, U: 0.389), a threshold of 0.05 and scanning across the full TAIR10 Arabidopsis genome. Sites were further filtered downstream to have a score of at least 4 – in the vast majority of cases corresponding to an exact match the (short) motif.

Generation of matched background regions: In order to account for sequence contexts specific to similar locations within the 3'UTR, each site from iCLIP, Nanopore and miCLIP was assigned a 'matched background' site, in a non-target gene, at the same relative location along the annotated genomic feature of the site (5'UTR, CDS or 3'UTR), according to a resolution of 10 bins per feature.

Motifs in the vicinity of ECT2 iCLIP/m$^6$A-Nanopore/ m$^6$A-miCLIP: motif distributions over distance centering on either iCLIP peaks or Nanopore peaks were generated using a custom R-script based on overlaps using *GenomicRanges* (Lawrence et al. 2013) At any given shift from the peakset, the raw number of overlaps of the motif (at any point) was calculated and normalized to give a motif count per 1000 peaks. To adjust for the potential for downstream regions over-shooting the end of the 3'UTR, at each given distance only sites that continue to overlap an annotated gene (Araport11) are counted. For the case where ECT2 iCLIP peaks are split by IDR status: a site is defined as IDR if it is in the 110 KDa set and greater than 10 nt away from a peak in the 55 kDa set, otherwise it is defined as non-IDR. For the case where Nanopore-defined m$^6$A sites are split according to the ECT2 target status of their containing gene, the target genes are defined according to ECT2-HT (roots+shoots), strict target genes are defined as target genes which are also in ECT2 iCLIP, and non-target genes are those with no evidence of ECT2 binding.

Gene body enrichment plots (metagene): to calculate motif enrichment over the gene body, motifs were first annotated a value according to their relative position within their relevant gene feature (5'UTR, CDS or 3'UTR). In order to account for over-representation of counts within the CDS, due to greater sequence coverage within transcript annotations, a random background set of 10 million positions were generated from the transcript annotation file and annotated in the same way as the motif locations to obtain an expected distribution of all positions over the gene features. This expected distribution was used to normalize the observed distribution of each motif and O/E values were plotted as a metagene plot over the gene. An enrichment of 1 suggests that the motif is neither over- or under- represented at that location.

Random forest analysis: called positions from either Nanopore $m^6A$ data (Parker et al. 2020) or ECT2 iCLIP were first reduced to remove redundant regions of multiple peaks within the same window, then paired with matched background sets (described above). Windows representing 'at' (+/- 10nt) the motif together with adjacent upstream 'up' and downstream 'down' windows of length 50nt (resulting in total window sizes of 120nt) around each position was annotated according to the number of each of the motifs overlapping (truncated at 10), and the final data set normalized. To create a held-out set, 1/5th of the peaks were removed from the set, and the other 4/5th were used to build a random forest model using gradient boosting (R package *gbm* (https://github.com/gbm-developers/gbm)), with settings specifying a shrinkage of 0.05, an interaction.depth of 6, cv.folds = 5 and n.trees = 2000. For each model ($m^6A$ Nanopore based or ECT2 iCLIP based) importance scores were extracted from the model and the top features were selected. The held-out data was further used to estimate the predictive score of the model by calculating the AUC (R package *pROC* (Robin et al. 2011). For each set-up, two further models were run – one involving the top 10 features from the full feature model, and one involving features from only DRACH and UNUNU (equating to 6 features in total), and AUC values calculated and compared to that of the full feature model.

### References (Supplemental Methods)

Arribas-Hernández L, Bressendorff S, Hansen MH, Poulsen C, Erdmann S, Brodersen P. 2018. An m6A-YTH Module Controls Developmental Timing and Morphogenesis in Arabidopsis. *Plant Cell* **30**: 952-967.

Arribas-Hernández L, Marchais A, Poulsen C, Haase B, Hauptmann J, Benes V, Meister G, Brodersen P. 2016. The slicer activity of ARGONAUTE1 is required specifically for the phasing, not production, of trans-acting short interfering RNAs in Arabidopsis. *Plant Cell* **28**: 1563-1580.

Bitinaite J, Nichols NM. 2009. DNA cloning and engineering by uracil excision. *Curr Protoc Mol Biol* **Chapter 3**: Unit 3 21.

Bolger AM, Lohse M, Usadel B. 2014. Trimmomatic: a flexible trimmer for Illumina sequence data. *Bioinformatics* **30**: 2114-2120.

Cheng C-Y, Krishnakumar V, Chan AP, Thibaud-Nissen F, Schobel S, Town CD. 2017. Araport11: a complete reannotation of the Arabidopsis thaliana reference genome. *Plant J* **89**: 789-804.

Dale RK, Pedersen BS, Quinlan AR. 2011. Pybedtools: a flexible Python library for manipulating genomic datasets and annotations. *Bioinformatics* **27**: 3423-3424.

Denyer T, Ma X, Klesen S, Scacchi E, Nieselt K, Timmermans MCP. 2019. Spatiotemporal Developmental Trajectories in the Arabidopsis Root Revealed Using High-Throughput Single-Cell RNA Sequencing. *Dev Cell* **48**: 840-852.e845.

Dobin A, Davis CA, Schlesinger F, Drenkow J, Zaleski C, Jha S, Batut P, Chaisson M, Gingeras TR. 2013. STAR: ultrafast universal RNA-seq aligner. *Bioinformatics* **29**: 15-21.

Grant CE, Bailey TL, Noble WS. 2011. FIMO: scanning for occurrences of a given motif. *Bioinformatics* **27**: 1017-1018.

Heinz S, Benner C, Spann N, Bertolino E, Lin YC, Laslo P, Cheng JX, Murre C, Singh H, Glass CK. 2010. Simple Combinations of Lineage-Determining Transcription Factors Prime cis-Regulatory Elements Required for Macrophage and B Cell Identities. *Mol Cell* **38**: 576-589.

Huppertz I, Attig J, D'Ambrogio A, Easton LE, Sibley CR, Sugimoto Y, Tajnik M, König J, Ule J. 2014. iCLIP: Protein–RNA interactions at nucleotide resolution. *Methods* **65**: 274-287.

Köster T, Staiger D. 2020. Plant Individual Nucleotide Resolution Cross-Linking and Immunoprecipitation to Characterize RNA-Protein Complexes. In *RNA Tagging: Methods and Protocols*, doi:10.1007/978-1-0716-0712-1_15 (ed. M Heinlein), pp. 255-267. Springer US, New York, NY.

Krakau S, Richard H, Marsico A. 2017. PureCLIP: capturing target-specific protein-RNA interaction footprints from single-nucleotide CLIP-seq data. *Genome biology* **18**: 240-240.

Lawrence M, Gentleman R, Carey V. 2009. rtracklayer: an R package for interfacing with genome browsers. *Bioinformatics* **25**: 1841-1842.

Lawrence M, Huber W, Pagès H, Aboyoun P, Carlson M, Gentleman R, Morgan MT, Carey VJ. 2013. Software for Computing and Annotating Genomic Ranges. *PLOS Computational Biology* **9**: e1003118.

Li H, Handsaker B, Wysoker A, Fennell T, Ruan J, Homer N, Marth G, Abecasis G, Durbin R, Genome Project Data Processing S. 2009. The Sequence Alignment/Map format and SAMtools. *Bioinformatics* **25**: 2078-2079.

Martin M. 2011. Cutadapt removes adapter sequences from high-throughput sequencing reads. *EMBnetjournal; Vol 17, No 1: Next Generation Sequencing Data AnalysisDO - 1014806/ej171200*.

Parker MT, Knop K, Sherwood AV, Schurch NJ, Mackinnon K, Gould PD, Hall AJW, Barton GJ, Simpson GG. 2020. Nanopore direct RNA sequencing maps the complexity of Arabidopsis mRNA processing and m6A modification. *eLife* **9**: e49658.

Patro R, Duggal G, Love MI, Irizarry RA, Kingsford C. 2017. Salmon provides fast and bias-aware quantification of transcript expression. *Nat Meth* **14**: 417-419.

Quinlan AR, Hall IM. 2010. BEDTools: a flexible suite of utilities for comparing genomic features. *Bioinformatics* **26**: 841-842.

Robin X, Turck N, Hainard A, Tiberti N, Lisacek F, Sanchez J-C, Müller M. 2011. pROC: an open-source package for R and S+ to analyze and compare ROC curves. *BMC Bioinformatics* **12**: 77.

Roehr JT, Dieterich C, Reinert K. 2017. Flexbar 3.0 – SIMD and multicore parallelization. *Bioinformatics* **33**: 2941-2942.

Shen L, Liang Z, Gu X, Chen Y, Teo Zhi Wei N, Hou X, Cai Weiling M, Dedon Peter C, Liu L, Yu H. 2016. N6-methyladenosine RNA modification regulates shoot stem cell fate in Arabidopsis. *Dev Cell* **38**: 186-200.

Wagih O. 2017. ggseqlogo: a versatile R package for drawing sequence logos. *Bioinformatics* **33**: 3645-3647.

Wei L-H, Song P, Wang Y, Lu Z, Tang Q, Yu Q, Xiao Y, Zhang X, Duan H-C, Jia G. 2018. The m6A Reader ECT2 Controls Trichome Morphology by Affecting mRNA Stability in Arabidopsis. *Plant Cell* **30**: 968-985.

Xu W, Rahman R, Rosbash M. 2018. Mechanistic implications of enhanced editing by a HyperTRIBE RNA-binding protein. *RNA* **24**: 173-182.
