## Supplemental Table 1 for "Principles of mRNA targeting via the Arabidopsis m^6^A-binding protein ECT2"

**Supplemental Table 1 - DNA Oligonucleotides (all sequences are 5' to 3')**

**Cloning**

| **Constructs** | **Primer pairs (fragments)** |
|---|---|
| *ECT2pro:ECT2-FLAG-ADAR-ECT2ter* (in pCAMBIA3300-U) | LA336-695 (*ECT2pro:ECT2*), LA696-615 (*FLAG-ADAR*), LA616-337 (*ECT2ter*) |
| *ECT2pro:FLAG-ADAR-ECT2ter* (in pCAMBIA3300-U) | LA336-697 (*ECT2pro*), LA698-615 (*FLAG-ADAR*), LA616-337 (*ECT2ter*) |
| *AGO1pro:FLAG-ADAR-AGO1ter* (in pCAMBIA3300-U) | MVUSER1-11 (*AGO1pro-FLAG*), MVUSER12-22 (*ADAR*), MVUSER23-6 (*AGO1ter*) |
| *FLAG-ADAR* (in pGEM-T Easy) | LA696-615 (*FLAG-ADAR*) |

**Primers/Oligonucleotides for USER-cloning**

| | |
|---|---|
| LA336.U-ECT2P.F | GGCTTAAUAAGCAACGAACCAAGGGAAGACG |
| LA337.ECT2T-U.R | GGTTTAAUAGGTTCTCTCGGCTTCTTTGAC |
| LA615.dADAR/ECT2T.R | AGTTATUCGGCAAGACCGAACTCGTC |
| LA616.dADAR/ECT2T.F | AATAACUAAGAGGATGGTGTCGCTC |
| LA695.ECT2/FLAG.R | ATCGCAACCAUTTGCCACCACATCG |
| LA696.ECT2/FLAG.F | ATGGTTGCGAUTACAAGGATGACGATGAC |
| LA697.ECT2P/FLAG.R | AATCCAUGAGAGGAGATTCGACAAACAAAG |
| LA698.ECT2P/FLAG.F | ATGGATUACAAGGATGACGATGAC |
| MVUSER1.F | GGCTTAAUCTATCCAAATTCCAAACCATACG |
| MVUSER6.R | GGTTTAAUGATTCTGTCGATTGCTTTGCTGG |
| MVUSER11.R | ATTGGACTGUACTTGTCATCGTCATCCTTG |
| MVUSER12.F | ACAGTCCAAUGGTGGTGCCACAG |
| MVUSER22.R | ACTGCGGCAGCUCATTCGGCAAGACCGAACTCG |
| MVUSER23.F | AGCTGCCGCAGUTGATTCACCCTCTATCTATCTTTATGACC |

**Primers for site-directed mutagenesis (QuickChange)**

| | |
|---|---|
| LA729.dADAR_E488Q_QC.F | CAAAATCGAGTCCGGTCAGGGGACGATTCCAG |
| LA730.dADAR_E488Q_QC.R | CTGGAATCGTCCCCTGACCGGACTCGATTTTG |

**Primers for detection of point mutations**

| | |
|---|---|
| LA660.dADAR_E488Q_CP(NlaIII).F | CGAAAACGACACTGGTGTTG |
| LA735.dADAR_E488Q_CP(NlaIII).R | GCTTTTCACTGGAATCGTCCCAT |
